# Volubility and vocal coordination under information asymmetry in captive common marmosets (*Callithrix jacchus*)

**DOI:** 10.64898/2026.01.24.701497

**Authors:** Monika Mircheva, Rahel K. Brügger, Judith M. Burkart

## Abstract

Volubility – the number of vocalizations per unit of time – is an understudied aspect of animal communication, potentially crucial in cooperatively breeding species that depend on coordinated behavior. Callitrichid monkeys such as common marmosets *(Callithrix jacchus)* are often characterized as highly vocal, yet variation in their calling rates remains poorly quantified. We recorded 69,538 vocalizations from 45 captive marmosets tested in dyads across six experimental conditions. Total call rates varied with condition but not with caller sex or status. Alertness call production depended on caller status: breeders showed context-appropriate responses to an ambiguous object, whereas helpers responded more generally. Food calls were tightly coupled to feeding rather than food availability, with males calling more than females. Dyadic analyses revealed call-type-specific covariation between subjects and partners, consistent with vocal contagion for alertness calls and recruitment for food calls. These patterns indicate flexible, call-type-specific volubility and information sharing beyond simple arousal contagion.

## Introduction

Vocal communication systems among animal species vary widely, mirroring the social complexity of each species^1,2^. For instance, animals living in larger groups have more extensive vocal repertoires^3,4^ and their vocalizations encode richer information^5^. Although group size is a good proxy, the quality of social interactions within a group may better illuminate the role of social complexity in communicative complexity. Cooperatively breeding species live in groups but also face unique social challenges compared to other group-living species. This breeding system is characterized by the involvement of individuals other than the parents in the care of offspring (i.e., alloparents), thereby making them highly interdependent. The need to engage cooperatively with multiple individuals regularly, often in quick succession, may necessitate precise and effective communication among agents to coordinate actions and share information. Indeed, cooperatively breeding birds have larger vocal repertoires than independently breeding ones, particularly in alarm and contact contexts^6^.

Beyond communicative complexity, in the sense of repertoire size, the propensity to produce vocalizations frequently may be an important and overlooked factor of vocal coordination. Volubility, defined as the number of vocalizations per unit time, could be especially relevant in cooperatively breeding systems, where frequent calling could help maintain contact, regulate group cohesion, and facilitate coordination. By contrast, in solitary-breeding species – where offspring care is primarily the mother’s responsibility and social coordination demands at this level are typically lower – calling rates may be comparatively reduced. Consistent with this idea, cooperative breeders such as meerkats show high and flexible volubility, e.g., by adjusting close-contact call rates during foraging to maintain group cohesion^7^.

In contrast, studies on independently breeding primates report relatively low call rates in the wild, including recent work on great apes in the context of mother-infant communication^8^, which aligns with earlier findings on volubility in adult chimpanzees, which produce only a few calls per hour^9^. Together, these results suggest that cooperative breeding may select for higher, context-sensitive vocal output due to the increased need for coordination of social interactions. Because humans exhibit key features of cooperative breeding^10^, studying the vocal communication of other cooperatively breeding species could offer valuable insights into the evolutionary conditions that support the emergence of a complex communication system, such as language, in humans.

Among primates, callitrichid monkeys are the only other cooperatively breeding species besides humans and demonstrate high levels of social tolerance and proactive prosociality^11–14^. Callitrichid caregivers readily provision infants without necessarily being prompted and sometimes even share food with other adult group members, with males tending to be particularly generous in this regard^15^. Callitrichids are also often considered highly voluble^16,17^, yet quantitative work on their volubility remains limited: to date, one study examined call rates in wild common marmosets in relation to diurnal and ontogenetic factors^18^, and one captive study measured time spent vocalizing as a function of social distance and its relationship to arousal^19^, but no study has systematically assessed how marmoset volubility varies across more specific contexts. Therefore, our understanding of potential variability in their volubility, contingent on contextual factors or the caller’s identity, remains limited.

The common marmoset is a widely used species for studying cooperative breeding systems and, therefore, a valuable model for investigating how the system-specific demands may shape vocal output. These small neotropical monkeys live in groups of three to fifteen individuals, typically comprising a dominant breeding pair, independent (i.e., helpers), and dependent (i.e., infant) offspring^20^. During early development, infants are typically carried on adults’ backs and passed between individuals, making efficient recruitment of potential carriers and coordination of infant transfers essential. Furthermore, sharing information about food resources and potential dangers in an environment with dense vegetation is crucial in a system as interdependent as cooperative breeding, and callitrichids’ rich vocal behavior makes the acoustic channel well-suited for such coordination. Common marmosets possess a vast vocal repertoire^21,22^ that is further expanded by the use of compound calls/call combinations^23,24^, providing a flexible communicative toolkit for navigating the challenges inherent in this highly interdependent social system.

Marmosets produce different types of contact calls based on the location of the intended receiver: the so-called “phee*”* calls are emitted as long-distance contact calls when individuals are visually separated and seek to reestablish contact with group members^22^. In contrast, “trill*”* calls serve as close-range contact calls when marmosets are in proximity to each other^25^. Closely related call types are “phee-peeps*”*, “trill-phees*”,* and “twitters*”*, whose functions are less well studied and thus less well understood, but they are likely to have a contact-seeking purpose as well. When employing contact calls, marmosets engage in vocal turn-taking and act as coupled oscillators^26^, closely resembling the way humans typically converse^27^. Furthermore, like other primates, marmosets produce functionally referential calls, including alarm and food calls^21,28^. Additionally, marmosets use a combination of three vocalization types (“tsik*”*, “ekk*”*, “ock*”*), singly or in sequence, with varying intensities, as mobbing calls or in situations of general uncertainty, frustration, anxiety, or fear^22,29^. Lastly, marmoset infants, like human infants, undergo a babbling phase and require vocal feedback from adults to refine their vocalizations^30–32^.

The objective of this study was to assess patterns of volubility of captive marmosets and to uncover why they vocalize so much. To do this, we recorded a large number of adult individuals of different sex-status classes. Dyads of individuals were placed in adjacent enclosures and could interact vocally across six experimental conditions. In the first condition, animals could see each other; in the next, visual contact between them was interrupted. In the following four counterbalanced conditions, visual contact remained interrupted, and a stimulus (an ambiguous object that could be interpreted as a potential threat or food) was presented to one dyad partner. These conditions created information asymmetry, and information available to only one individual could potentially be shared with the unaware dyad partner. Although our knowledge of marmoset volubility is limited, we can nevertheless predict patterns based on the social roles of different status classes (breeder vs. helper) and sex classes in marmosets’ everyday behavior, which may be linked to who shares information with whom in specific contexts.

Firstly, we expected total volubility to increase when marmosets were separated visually from each other. Visual separation would result in higher volubility due to increased contact-seeking and information sharing under the respective conditions. Specifically, as contact calls are typically produced when animals are separated from their group members^19,22^, we expected an increase in contact-seeking vocalizations when visual contact between partners was obstructed. Moreover, we may observe an effect of age on contact-seeking behavior, with younger individuals being more insecure when separated from group members and keener to maintain contact with their partner. Secondly, marmosets use “alarm calls” and “tsik”/”ekk”/”ock” calls when facing perceived danger, but also use the latter in situations of general uncertainty, frustration, and anxiety^29^. Therefore, we expected that individuals would produce these alertness vocalizations at a higher rate when confronted with an ambiguous object they might consider dangerous, or when visually separated from their groupmates, which may increase arousal. In this case, we again anticipated an effect of age, with younger individuals lacking experience with novel, ambiguous stimuli and environments and producing more “tsik”/”ekk”/”ock” and “alarm calls”. Thirdly, food-associated calls are closely linked to the presence and consumption of food^28,33^; therefore, we predicted that marmosets would produce these calls when food was available. Additionally, we expected that males^34^ might be more inclined to share information about the discovered food and produce food calls to recruit other group members.

Lastly, we were interested in how individual volubility in situations of information asymmetry – such as having access to ambiguous stimuli or food while the partner is unaware of them – might affect the interaction partner’s volubility. In conditions with ambiguous stimuli, we expected that the production of alertness calls by the individual interacting with the stimulus would trigger an increase in the partner’s alertness calls, as a social contagion effect^35^. Likewise, it might prompt increased contact-seeking to signal that the partner is still nearby and de-escalate the situation. Furthermore, when one individual has access to food while the other does not, the individual without access may increase alertness vocalizations as a manifestation of frustration over the inequity in access to the food source, i.e., inequity aversion^36^. Sharing information about food – such as producing food calls – might also increase contact-seeking in the partner whose visual access is obstructed, signaling their presence and willingness to join in feeding.

## Results

### Total call rate across conditions

We recorded a total of 69,538 vocalizations (Table 2) and first investigated how total volubility, encompassing all call types, varies across different experimental conditions (model 1). The reduced zero-inflated negative binomial GLMM provided a significantly better fit than the null model (likelihood ratio test, *N*_total_ = 1380, *N*_individual_ = 45, *N*_group_ = 14, χ^2^ _(17)_ = 427.84, *p* < .001; Supplementary Table S4) and revealed a significant effect of experimental condition on total volubility (χ^2^ _(5)_ = 96.04, *p* < .001, Supplementary Table S5). We found a significant difference between the visual condition and all other experimental conditions (odds ratio = 1.42, 95% CI [1.18; 1.71], *z* = 3.68, *p* < .001; Figure 1A&B), with individuals having 42% higher call rates when visual contact was obstructed. However, we did not detect an increase in calling in the stimuli conditions compared to the non-visual neutral condition (odds ratio = 0.95, 95% CI [0.81; 1.11], z = −0.69, p = 0.49; Figure 1A&B). Additionally, the model predicted a significant increase in calling when individuals interacted directly with stimuli (ambiguous/food-for-self) compared to when they were passive partners (ambiguous/food-for-partner, odds ratio = 2.25, 95% CI [1.67; 3.04], *z* = 5.30, *p* < .001; Figure 1A&B), with the former having 125% higher call rates. Furthermore, we also observed a significant difference between the two active interaction conditions (ambiguous-for-self vs food-for-self; odds ratio = 2.37, 95% CI [1.90; 2.95], *z* = 7.69, *p* < .001; Figure 1A&B). The model estimated that individuals have roughly a 2.4-fold higher call rate when they encounter food than when they discover the ambiguous object. There was a trend for individuals to call more in the food condition than in the ambiguous condition when they were the naïve partner (ambiguous/food-for-partner; odds ratio = 1.22, 95% CI [0.99; 1.49], z = 1.90, p = 0.057, Figure 1A&B), with marmosets having 22% higher calling rates when their partner, but not they themselves, was encountering food. We found an effect of session number on calling behavior (odds ratio = 0.93 per session, 95% CI [0.90; 0.97], χ^2^ _(1)_ = 13.17, *z* = –3.63, *p* < .001; Supplementary Table S5, Supplementary Figure S3A), indicating a 7% decrease in calling rate per test session. Individual features such as caller sex, status, or age did not affect overall call rates (Supplementary Table S5).

**Figure 1:**
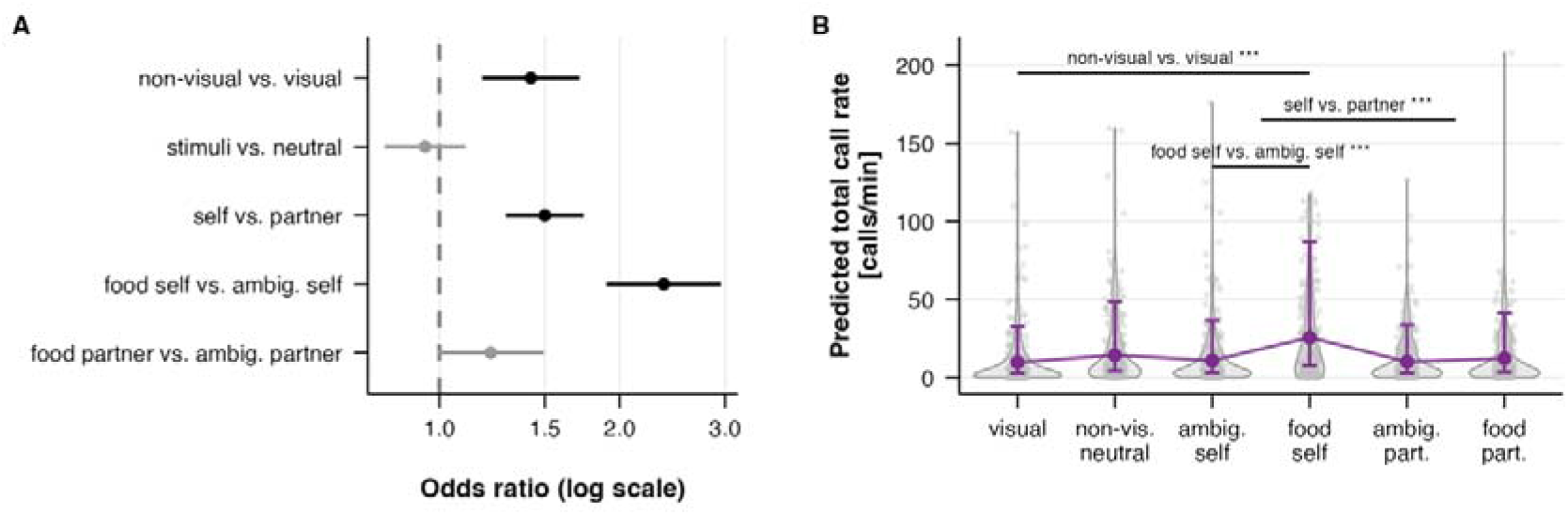
Effects of experimental condition on total call rate*. **(A)** Odds ratios with 95% confidence intervals for a priori planned contrasts testing: whether call rates differed between all non-visual conditions and the visual condition (non-visual vs. visual); whether stimulus presence affected call rates under visual separation (stimuli vs. neutral); whether information asymmetry between dyad partners influenced call rates (self vs. partner); and whether call rates differed between food and ambiguous objects when the subject (food vs. ambig. self) or their partner (food vs. ambig. partner) had access to the stimulus. Significant contrasts (p < .05) are shown in black, non-significant contrasts in grey. The dashed vertical line indicates an odds ratio of 1 (no effect). **(B)** Model-predicted call rates, including all call types across experimental conditions: when dyad partners could see each other (visual), when visual contact was obstructed without additional stimuli (non-vis. neutral), when visual contact was obstructed, and the subject could interact with an ambiguous object or food (ambig./food self), and when their partner could interact with the stimulus (ambig./food part.). Points and error bars represent model-estimated means and 95% confidence intervals, respectively. Violin plots show the distribution of observed call rates. Dots indicate individual observations. Brackets indicate significant contrasts. *Panels A and B are equivalent

### Contact call rates

We furthermore examined how contact call rates varied across experimental conditions (model 2). The reduced zero-inflated negative binomial model significantly outperformed the null model (likelihood ratio test, *N*_total_ = 1380, *N*_individual_ = 45, *N*_group_ = 14, X*^2^*_(16)_ = 351.86, *p* < .001; Supplementary Table S4). However, the experimental condition did not significantly affect contact call rates (χ^2^_(5)_ = 5.17, *p* = 0.40; Supplementary Table S5), indicating that none of the planned contrasts reached significance. As with the total volubility model including all call types, we found a significant effect of test session on subjects’ call rate (odds ratio = 0.95 per session, 95% CI [0.93; 0.99], *z* = –2.85, X*^2^*_(1)_ = 8.15, *p* = 0.004; Supplementary Figure S3B), indicating a 5% decrease in call rate with each subsequent test session. We found no significant effects of individual features such as caller sex, status, or age on contact-seeking call rates (Supplementary Table S5). However, we observed a trend suggesting that caller status influenced contact call rates (X*^2^*_(1)_ = 3.59, *p* = 0.06), with helpers showing marginally higher rates than breeders (odds ratio = 1.57, 95% CI [0.98; 2.51], z = 1.90).

### Alertness call rates

For alertness calls, the reduced model 3A significantly outperformed the null model (likelihood ratio test, *N*_total_ = 1371, *N*_individual_ = 45, *N*_group_ = 14, X*^2^*_(23)_ = 241.46, *p* < .001, Supplementary Table S4) and revealed a significant interaction effect between experimental condition and caller status (χ^2^_(6)_ = 32.85, *p* < .001; Supplementary Table S5). Given this significant interaction, we examined the condition contrasts separately for breeders and helpers. For breeders, alertness call rates were significantly higher in stimulus conditions compared to the neutral non-visual condition (odds ratio = 0.51, 95% CI [0.33; 0.80], z = −2.93, *p* = 0.003), and within the stimulus conditions, significantly higher in food than in ambiguous object conditions (odds ratio = 0.37, 95% CI [0.24; 0.57], z = −4.43, *p* <.001). Breeders who encountered the ambiguous object themselves produced fewer alertness calls than when their partner encountered it (odds ratio = 0.52, 95% CI [0.27; 0.99], z = −1.91, *p* = 0.048). Critically, breeders who approached the ambiguous object showed approximately 12-fold higher alertness call rates compared to those who did not approach (odds ratio = 12.22, 95% CI [4.50; 33.17], z = 4.91, *p* <.001). Helpers followed a different pattern: alertness call rates did not differ between non-visual and stimulus conditions (odds ratio = 0.82, 95% CI [0.58; 1.16], z = −1.13, *p* = 0.258), nor between food and ambiguous object conditions (odds ratio = 1.41, 95% CI [0.97; 2.05], z = 1.81, *p* = 0.071). Interestingly, opposite to the breeder pattern, helpers who encountered the ambiguous object produced higher alertness call rates than then their partner did (odds ratio = 1.70, 95% CI [1.01; 2.86], z = 2.00, *p* = 0.046) regardless of whether they actually approached the object or not (odds ratio = 0.77, 95% CI [0.33; 1.77], z = −0.62, *p* = 0.534). The breeder-helper differences were statistically significant for the pooled ambiguous-vs-food contrast (odds ratio = 0.26, 95% CI [0.15; 0.46], z = −4.57, *p* <.001), the ambiguous-for-self-vs-ambiguous-for-partner contrast (odds ratio = 0.31, 95% CI [0.13; 0.70], z = −2.81, *p* = 0.005) and the approach effect (odds ratio = 15.95, 95% CI [4.34; 58.59], z = 4.17, *p* <.001).

**Figure 2:**
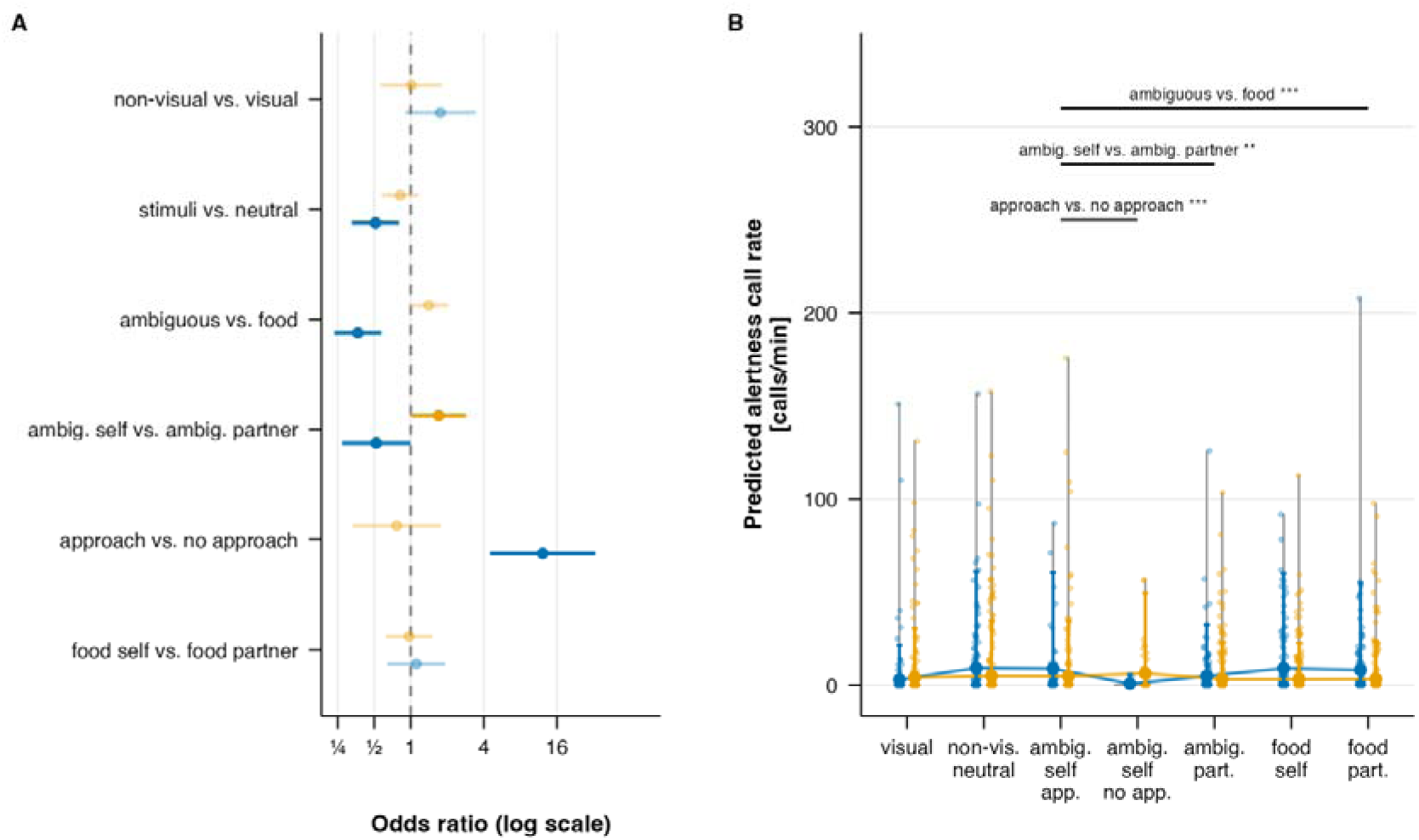
Effects of experimental condition on alertness call rates*. **(A)** Odds ratios with 95% confidence intervals for a priori planned contrasts testing whether: alertness call rates differed between all non-visual conditions and the visual condition (non-visual vs. visual); stimulus presence affected alertness calls under visual separation (stimuli vs. neutral); alertness call rates differed between ambiguous object and food conditions, pooled across for-self and for-partner (ambiguous vs. food); information asymmetry influenced alertness calling in ambiguous object conditions (ambig. self vs. ambig. partner); direct interaction with the ambiguous object affected alertness calling (approach vs. no approach); information asymmetry affected alertness call rates in food condtions (food self vs. food partner). Significant contrasts (p<.05) are shown at full color saturation; non-significant contrasts are faded. The dashed line indicates an odds ratio of 1 (no effect). **(B)** Model-predicted alertness call rates across experimental conditions: when dyad partners could see each other (visual), when visual contact was obstructed without additional stimuli (non-vis. neutral), when the subject encountered an ambiguous object, with or without approaching it (ambig. self app./no app.), when their partner encountered the ambiguous object (ambig. part.), and when the subject or their partner had access to food (food self/part.). Points and error bars represent model-estimated means and 95% confidence intervals, respectively. Violin plots show the distribution of observed call rates. Dots indicate individual observations. Brackets indicate significant between-status differences in the corresponding contrasts. In both panels, blue and orange represent breeders and helpers, respectively. *Panels A and B are equivalent

The model also revealed a significant effect of caller age (odds ratio = 0.98 per month, 95% CI [0.97; 1.00], *z* = –2.29, *p* = 0.022; Supplementary Table S5) on alertness call production across conditions, with younger individuals calling more frequently than older ones (Figure 3). Animals showed a 21% decrease in call rate per year of age. Additionally, as with contact calls, we observed a habituation effect (odds ratio = 0.92 per session, 95% CI [0.86; 0.98], z = –2.15, *p* = 0.015; Supplementary Table S5, Supplementary Figure S3C), with alertness call rates decreasing by about 8% per test session.

**Figure 3:**
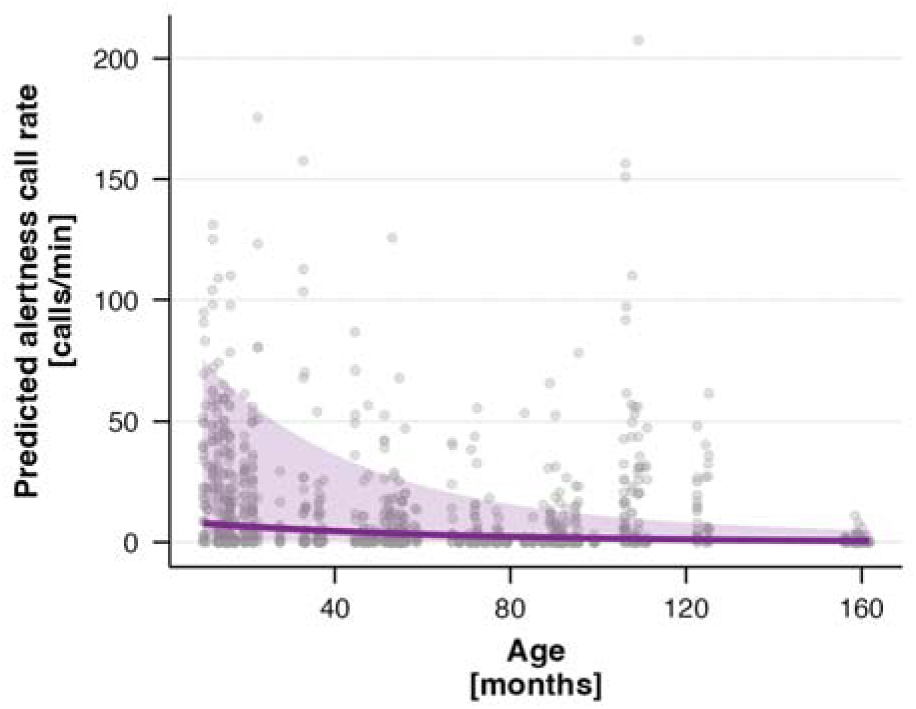
Effect of age on alertness call rates. The purple line and shaded area indicate the model-predicted trend and 95% confidence intervals, respectively. Dots represent individual observations.

### Partner responses to alertness calls

The next model examined how a subject’s alertness calls affect the naïve partner’s production of alertness calls in the ambiguous object condition (model 3B). In this condition, only the subject had access to the ambiguous object, while the partner had no visual access to it nor to the subject (information asymmetry) and could only rely on the subject’s vocal behavior. The reduced model provided a significantly better fit than the null model (likelihood ratio test, *N*_total_ = 221, *N*_individual_ = 45, *N*_group_ = 14, χ^2^_(7)_ = 60.49, *p* < .001, Supplementary Table S4) and revealed that subjects’ alertness calls significantly predicted their partner’s alertness calling (χ^2^_(1)_ = 41.04, *p* <.001). Each doubling of calls a subject produced was associated with a 1.4-fold increase in alertness calls produced by the partner (odds ratio = 1.68 per 10 calls, 95% CI [1.43; 1.97], z = 6.41), with the strongest contagion at low subject call rates and diminishing returns at higher rates (Figure 4). No other predictor showed a significant effect.

**Figure 4:**
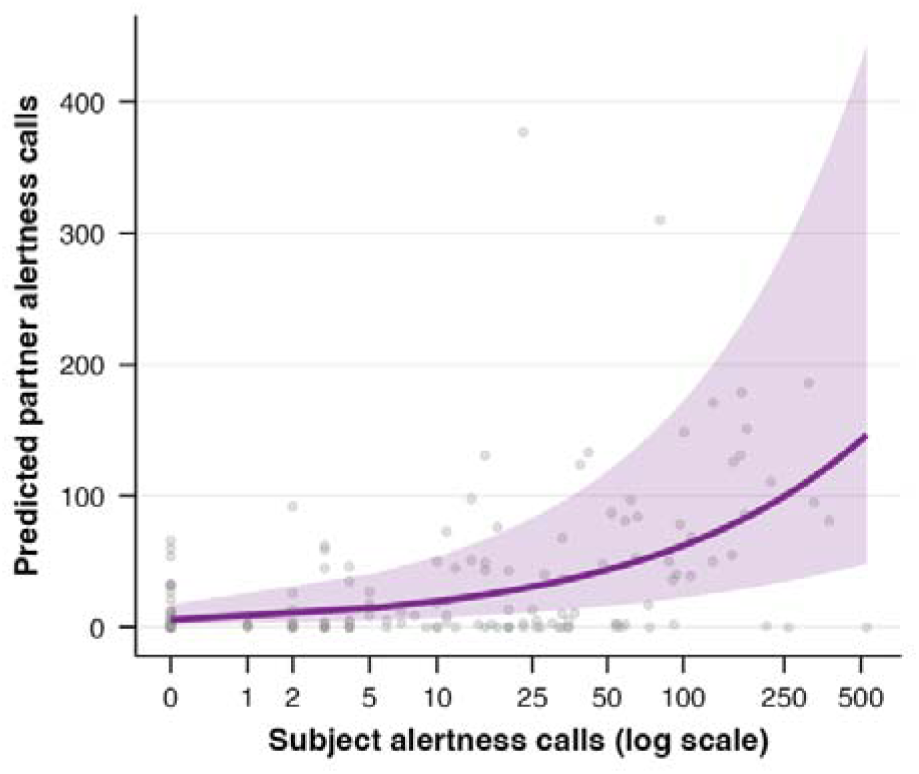
Vocal contagion of alertness calls in the ambiguous-for-self condition. The purple line and shaded area indicate the model-predicted partner alertness call rate as a function of the subject’s alertness call rate, with the 95% confidence interval. Dots represent individual observations.

Furthermore, to test whether naïve partners might respond to the subject’s alertness calls by producing more contact calls – potentially serving a de-escalation or proximity-signaling function – we examined whether subject’s alertness calls were positively associated with partner’s contact call production (model 3C). The reduced model fit the data significantly better than the null model (likelihood ratio test, *N*_total_ = 221, *N*_individual_ = 45, *N*_group_ = 14, χ^2^_(9)_ = 27.89, *p* < .001, Supplementary Table S4). However, we found no significant correlation between subject’s alertness calls and their partner’s contact calls (χ^2^_(1)_ = 1.05, *p* = 0.31).

### Food call rates

In the second information-asymmetry condition, preferred food became available to the subjects but not to their partners. To capture both the condition-level effects and the subject’s manner of interaction with the food in a single analysis, we modeled food call rates as a function of a nested variable with four levels: approach with feeding, approach without feeding, no approach (all 3 within the food-for-self condition), and food for partner condition (model 4A). The reduced model fit the data significantly better than the null model (likelihood ratio test, *N*_total_ = 426, *N*_individual_ = 45, *N*_group_ = 14, χ^2^_(9)_ = 209.25, *p* < .001, Supplementary Table S4) and revealed a significant effect of the nested condition variable (χ^2^_(3)_ = 137.75, *p* < .001; Supplementary Table S5). The overall comparison between food-for-self and food-for-partner did not reach significance (odds ratio = 1.17, 95% CI [0.90; 1.50], z = 1.18, *p* = 0.24). However, the dominant effect was driven by individuals who approached and fed on the food, who produced dramatically higher food call rates compared to all other categories (odds ratio = 6.44, 95% CI [4.37; 9.49], z = 9.40, *p* < .001; Figure 5A&B). Interestingly, some individuals who approached and ate the food made no food calls at all (Figure 5B). In contrast, approaching the food without consuming it did not significantly increase food call rates relative to not approaching it at all (odds ratio = 1.32, 95% CI [0.58; 3.01], z = 0.67, *p* = 0.506). Furthermore, the caller’s sex had a significant effect on food call rates (χ^2^_(1)_ = 5.20, *p* = 0.023; Supplementary Table S5), with males producing higher rates than females (odds ratio = 2.7, 95% CI [1.15; 6.35], z = 2.28; Figure 5C). There were no effects of status or age, nor any indication of habituation effects (Supplementary Table S4).

**Figure 5:**
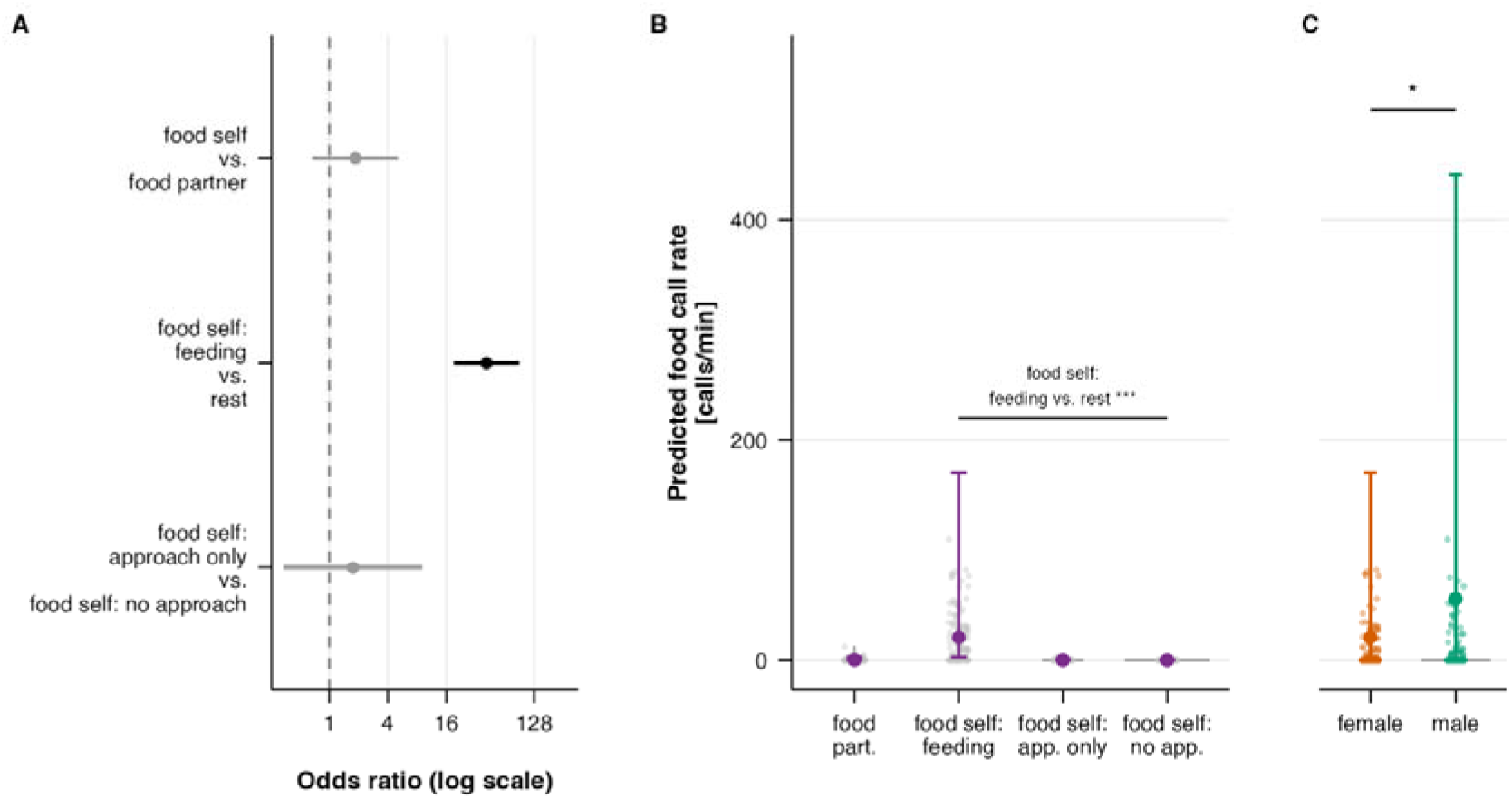
Effects of experimental condition, manner of interaction with food, and caller sex on food call rate*. **(A)** Odds ratios with 95% confidence intervals for a priori planned contrasts testing whether: food call rates differed between food-for-self and food-for-partner conditions (food self vs. food partner); food consumption affected food calling compared to only approaching the food and not approaching at all (food self: feeding vs. rest); approaching without feeding differed from not approaching at all (food self: approach only vs. food self: no approach). Significant contrasts (p < .05) are shown in black; non-significant contrasts in grey. The dashed vertical line indicates an odds ratio of 1 (no effect). **(B)** Model-predicted food call rates across the food conditions: with food self split in when the subject approached and fed on the food (feeding), when the subject approached but did not feed (app. only), and when the subject did not approach the food at all (no app.); and when the dyad partner had access to the food (food part.). **(C)** Model-predicted food call rates by caller sex. In panels B and C, points and error bars represent model-estimated means and 95% confidence intervals, respectively. Violin plots show the distribution of observed call rates. Dots indicate individual observations. Brackets indicate significant contrasts. *Panels A and B are equivalent

**Figure 6:**
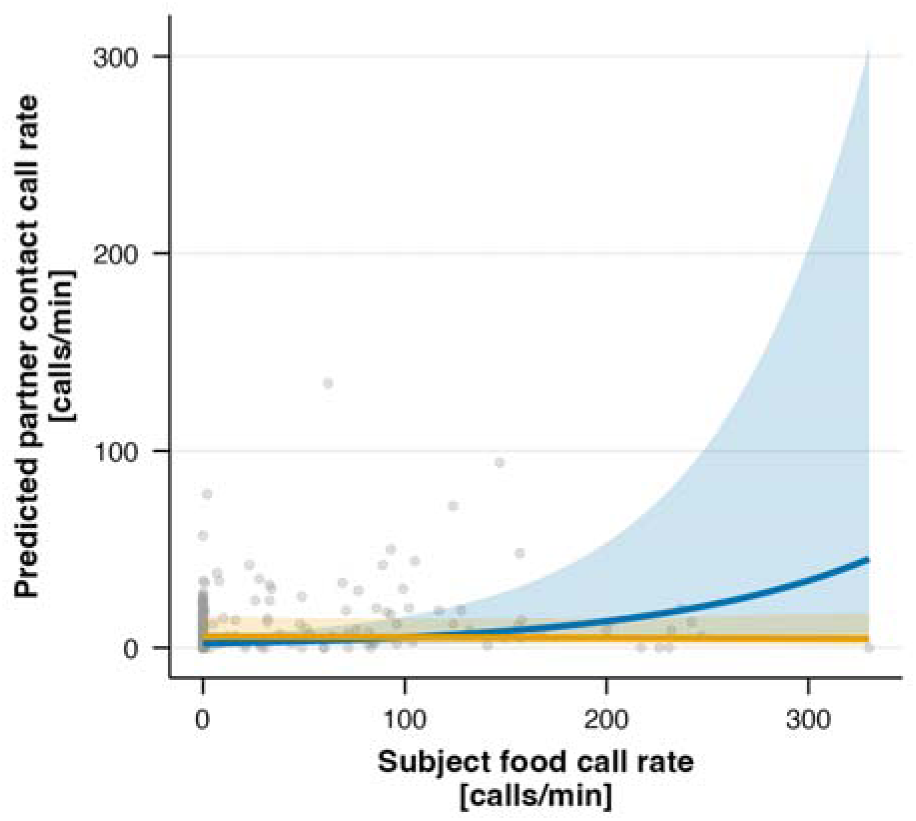
Partner contact calls as a function of subject food calls in the food-for-self condition. Lines and shaded areas indicate model-predicted trends and 95% confidence intervals for breeders (blue) and helpers (orange) as focal subjects, respectively. Dots represent individual observations.

### Partner responses to food calls

Next, we examined how the subjects’ food calls influenced their naïve partners’ vocal behavior, as they had no direct access to the food and could only perceive the subject’s vocal cues. The reduced model examining how a caller’s food calls influence their partner’s contact-seeking vocalizations provided a significantly better fit than the null model (model 4B; likelihood ratio test, *N*_total_ = 223, *N*_individual_ = 45, *N*_group_ = 14, χ^2^_(9)_ = 48.75, *p* < .001, Supplementary Table S4) and revealed a significant relationship between the two variables, depending on the caller’s status (χ^2^_(1)_ = 9.63, *p* = 0.002; Supplementary Table S5). Model predictions showed that increases in breeders’ food calls were associated with a significant rise in their partners’ contact calls (trend = 0.08, 95% CI [0.03; 0.12]). This effect was absent in helper food calls (trend = −0.01, 95% CI [-0.06; 0.04]) and the difference between status classes was significant (estimate = 0.09, z = 2.36, *p* = 0.02).

We also tested whether subjects’ food calls covary with their partners’ alertness calls (model 4C), examining whether partners might react to food-associated vocal signals with heightened alertness or annoyance when they cannot reach the food themselves. However, the model did not significantly improve upon the null model (likelihood ratio test, *N*_total_ = 223, *N*_individual_ = 45, *N*_group_ = 14, χ^2^_(7)_ = 8.34, *p* = 0.304, Supplementary Table S4), providing no evidence towards a potential inequity aversion reaction by the partner.

## Discussion

### Total call rate across conditions

In this study, we systematically quantified and described the volubility of captive common marmosets across various experimentally manipulated contexts and individual characteristics. We did not find apparent differences in total volubility encompassing all call types between sex or status classes; instead, volubility varied primarily with experimental condition. As expected, call rates were generally higher during visual occlusion, highlighting increased reliance on the acoustic channel when visual communication with social partners is unavailable. This pattern is consistent with Liao et al. (2018)^19^, who reported that marmosets spend more time vocalizing when separated from social partners, and aligns with the broader idea that vocal signaling becomes especially important when other communication modalities are constrained^37^. Our supplementary trend analysis confirms that this pattern is not driven by a linear temporal decline within the session (Supplementary Table S6). Such reliance on acoustic communication is particularly relevant for common marmosets, who inhabit dense vegetation where visual contact is often limited, making vocal interactions, often over long distances, more critical^25,38,39^.

Marmosets increased their call rates when actively engaging with the provided stimuli (ambiguous objects or food) compared to when they were the ignorant party, i.e., when their dyad partner was exposed to the stimuli. This shows that direct interaction with ecologically relevant stimuli elicits vocal reactions that can function to inform or recruit an uninformed partner. Such context-dependent increases in volubility are consistent with a communicative role for these calls rather than purely arousal-driven vocal output due to separation from social partners^40^. This interpretation is further supported by results from the subject-partner vocal dynamics, when taking call type categories into account, as detailed below.

### Contact call rates

Contact-seeking vocalizations (i.e., phee, trill, trill-phee, and twitter) did not show the predicted condition effects. We expected contact calling to increase during visual separation from the partner, but we did not find a significant effect of the experimental condition. The absence of a condition effect is consistent with our supplementary trend analysis, which also yielded no significant condition or condition order effect (Supplementary Table S6), suggesting that the null result is not an artifact of temporal position within the session. These results appear at odds with Liao et al. (2018)^19^, who reported greater vocal output during visual separation in a dataset dominated by contact calls. However, it is likely that the fixed position of the visual condition at the start of each session, combined with the known exponential decline in phee call production over time^41^, may have masked a genuine increase in contact calling during visual separation, which would have been detectable had condition order been fully randomized as in Liao et al. In our experiment, we intentionally placed the visual condition first to ensure animals were aware of who their partner was in all subsequent information-asymmetry conditions under visual separation. Contact calling also decreased over test sessions, further supporting the notion that prolonged and repeated exposure leads to reduced vocal output^25,42^.

We, furthermore, predicted that contact call rates would be higher when individuals were exposed to ecologically relevant stimuli (an ambiguous object or food), consistent with contact calls serving a recruiting function. Specifically, increased contact calling in food conditions could signal the presence of food and encourage group members to join in at the feeding site, while increased contact-seeking in the ambiguous object condition could recruit partners for a coordinated response to a potential threat. Both would align with marmosets’ cooperative breeding system, in which advertising resources and coordinating responses to potential dangers are essential for group cohesion and survival^14^. However, we found no significant difference in contact call rates across conditions, and thus no support for this hypothesis. Furthermore, we predicted that younger individuals would show higher contact-seeking. We found no age effect on contact call rates but a marginal trend for status: helpers tended to produce contact calls at higher rates than breeders across conditions of visual separation. It may thus be that helpers (who also tend to be younger in our colony and can be reproductively suppressed in their natal group^43^) use separation as an opportunity to produce calls with functions beyond contact maintenance with the natal group (e.g., mate attraction or territorial advertisement). Because our analyses grouped vocalizations into broad functional categories, disentangling these possibilities will require further acoustic analyses to distinguish contact calls used to regain contact from those serving other functions^44^ and testing whether contact calls co-occurring with food calls have distinct acoustic structures consistent with recruitment, as has recently been suggested (Briefer et al. subm).

### Alertness call rates

As expected, we found that alertness call production varied with experimental condition, depending on the caller’s status, with marked differences between breeders and helpers. Breeders produced more alertness calls when *their partner* encountered the ambiguous object than when they themselves did. They likely interpreted the ambiguous object as less threatening than helpers did, yet they still responded strongly to partners’ alertness calls because they could not know the reason for their partners’ alertness. Consequently, when breeders were presented with the ambiguous object, their alertness call rate rose only when they approached it, compared to when they did not, the latter arguably occurring when they were uncertain about it. We cannot distinguish whether breeders called more when approaching to check out the object about which they were uncertain, or if the approach itself was already a part of an unfolding mobbing response that began with calling. Intriguingly, breeders also produced significantly more alertness calls in both food conditions than in ambiguous-object conditions, whereas helpers did not. While we can’t provide a fully satisfactory explanation, one possibility is that some individuals produced these calls in protest of their partner having access to food while they did not. However, we found no association between subject food calls and partner alertness calls (model 4C), which speaks against a frustration-driven interpretation. A more plausible alternative arises from the anecdotal observation that many individuals produced tsik-ekk calls as they approached the food. This is consistent with Bosshard et al. (2024)^24^, who showed that marmoset call sequences combine food-related and mobbing-related calls into recurring combinations, suggesting that the boundary between these contextual categories is less strict than previously thought. The live, moving mealworms used here may also have contributed to elevated arousal during the food encounter, blurring the line between food-related and alertness-related vocalizations.

Among helpers, alertness call rates did not vary significantly across conditions, but they produced more alertness calls when they encountered the ambiguous object themselves, a pattern that differed significantly from breeders, in whom the direction was reversed. Unlike breeders, this increase in helpers was independent of whether they approached the object, and the approach effect itself differed significantly between the two status classes. Helpers appear to be more reactive to the ambiguous stimuli in general, producing alertness calls regardless of whether they engage with the object. Consistent with this, we found a robust age effect on alertness call production, with younger individuals calling at higher rates. Because helpers in our sample are typically younger and less experienced, they may be more easily aroused by unfamiliar stimuli and perceive them as more threatening even from a distance, especially when combined with encountering them on their own, outside their home enclosure with only one dyad partner present. Consistent with this interpretation, marmosets exposed to unfamiliar environments have been shown to produce more “tsik-ekk*”* calls, i.e., alertness calls, likely reflecting heightened general uncertainty and anxiety^29^.

### Partner responses to alertness calls

We found a clear positive association between focal subject and partner alertness call rates in the ambiguous object condition, consistent with vocal contagion^45^. By contrast, we found no such relationship between the focal individual’s alertness calls and their partner’s contact calls (model 3C), and thus no evidence that partners attempted to de-escalate the situation, in other words, calm a distressed conspecific by signaling proximity. This makes sense biologically because ignoring a partner’s alertness calls without knowing the reason for them might lead to overlooking a real danger or predator. Alertness calls, such as “tsik*”* calls, are often associated with mobbing and mobbing-like anti-predator contexts^22,46,47^. From a functional perspective, matching a partner’s arousal state and joining in a coordinated response would indeed be more adaptive than producing contact calls – particularly when an individual cannot directly assess the stimulus but can use their partner’s vocalizations as an indicator of potential threat. Finally, alertness call rates by focal subjects declined remarkably over test sessions, likely reflecting habituation to the experimental conditions and repeated exposure to the ambiguous object, thereby reducing novelty-related arousal over time.

### Food call rates

Overall, focal subjects did not produce more food calls in the food-for-self vs. food-for-partner condition. Instead, food call production was mostly driven by food consumption: marmosets called most when approaching and then feeding on the food, whereas food presence or approaching it without feeding elicited little to no calling. This suggests that marmosets do not necessarily advertise the presence of food even though they are aware of it. Notably, food calls were the only call type that did not show a clear habituation effect across sessions, suggesting that food remained a highly salient stimulus throughout the experiments^42^.

Food availability and consumption were arguably associated with elevated arousal^48^, and variation in food motivation may have contributed to individual differences in food calling. In this case, food calling would directly reflect arousal. However, several individuals consistently refrained from producing food calls despite readily and eagerly consuming the food, suggesting that food calling is not simply an automatic by-product of arousal. Instead, some individuals may have voluntarily suppressed it contingent on additional factors such as social context or individual prosocial tendencies. Consistent with this idea, and stronger prosocial tendencies in males (particularly in helpers^34^), males produced more food calls than females. The absence of a status effect on food calling among males may reflect individual differences in prosociality, as well as the relationships among dyadic partners, which should be addressed in future studies.

### Partner responses to food calls

Subjects seldom produced food calls when they were the passive partner (food-for-partner condition), even when their dyad partner was food calling. This shows that in contrast to alertness calls, food calls are not contagious. This is adaptive because they are thus only given at the location where food actually is, which facilitates localizing a food source by other group members.

Rather than eliciting food calls contagiously, food calls produced by breeders elicited increased contact calling from dyadic partners (regardless of that partner’s status). This is consistent with the notion that breeder calls might be a particularly salient signal for initiating and coordinating interaction, given that breeders are dominants within and build the core of a social group^20^. Notably, we did not observe an increase in partners’ alertness calls when the subject produced food calls, providing no evidence that partner vocal behavior reflected frustration at unequal access to food and thus no support for inequity aversion, at least in this context. This finding is consistent with marmosets’ increased prosociality^11,34^.

### Limitations and conclusions

It is important to note that because our analyses grouped vocalizations into broader functional categories, we cannot determine whether finer-grained call-type usage differs across sex-status classes or dyad compositions (e.g., breeder-breeder vs breeder-helper). Although we recorded a large number of vocalizations, some call types occurred much less frequently than others, limiting statistical power for more detailed comparisons at the call-type level. We also note that our findings come from captive animals tested in an experimental setting that differs in important ways from natural conditions. In our experiment, subjects could not freely approach group members, and the dyadic setup isolated them from the rest of the social group during test sessions. While this controlled setting allowed us to create situations of information asymmetry in alertness and food contexts in specific dyads, and isolate vocal behavior, the extent to which these patterns generalize to wild marmosets remains an open question and motivates future fieldwork.

Overall, the present study demonstrates that marmoset volubility is variable and heavily influenced by social context and individual traits. Vocalization rate across call type categories is affected by different factors, suggesting that volubility is not a uniform trait but rather more flexible and complex. The experimentally induced information-asymmetry conditions are particularly informative regarding the factors shaping vocal output, and to what extent it reflects arousal and information sharing.

Individuals increased their calling rates when directly engaged with salient stimuli, consistent with elevated arousal-driven vocal output, in particular for alertness calls: Alertness call rates correlated negatively with age and thus experience, and repeated exposure to (and thus experience with) the ambiguous object led to a decrease. Moreover, they positively covaried between subjects and partners in the ambiguous object condition, consistent with vocal and emotional contagion that can lead to coordinated threat responses, such as joining in mobbing without directly perceiving the potential threat. For food calls, however, the dyadic analyses revealed call-type-specific associations between subject and partner call rates that go beyond what a simple arousal account could predict. Food calls given by breeders encountering the food were positively correlated with partners emitting more contact calls. These covariation patterns indicate the presence of functionally specific social influences on vocal output rather than generalized arousal contagion, though the directionality of these relationships cannot be determined from the present analyses. Individual variation in food calling, including more calling by males (who tend to be more prosocial^34^) and individuals that readily and eagerly consumed food without vocalizing, further suggests that factors beyond arousal state alone, such as individual differences in prosocial tendencies or food motivation and food competition (in particular, females^49^), appear to modulate vocal output.

Together, our findings support the idea that volubility may play an important role in a cooperative breeding system, where recruitment, information sharing, and rapid coordination may be particularly beneficial. A key next step is to compare which within-species patterns of context-dependent volubility are specific to cooperatively breeding species by comparing them to independently breeding ones. Once established, this information will help delineate the role that cooperative breeding may have played during human evolution. Humans are the only cooperative breeders among great apes; they are characterized not only by linguistic complexity but also by an exceptional degree of vocal output^50^, using language to express complex ideas^51^, coordinate joint action^52^, and sustain large-scale cooperation^53^. Higher volubility and information sharing, once our ancestors started to engage in extensive allomaternal care and cooperative breeding, may well have been a strong initial impetus towards the evolution of our unique communication system, language^54,55^.

## Methods

### Study subjects

We recorded 45 captive common marmosets, including 22 females, from eight family groups and six couples (see Supplementary Table S1 for details). Family groups consisted of a breeding pair and up to two helpers, while couples included only a breeding pair without offspring. The animals were housed in indoor enclosures (2.7 x 1.8 x 2.4 m) with outdoor access (3.2 x 1.8 x 2.4 m) available when weather conditions permitted. Marmosets received a vitamin supplement mash in the morning and a variety of vegetables at lunch. An afternoon snack (e.g., egg, cricket, or raisin) was provided along with mealworms and gum arabic. Fresh water was available ad libitum.

### Experimental setup

The experiments were conducted in an experimental room equipped with two enclosures (1 x 1.8 x 2 m; Figure 7A). Marmosets could access these enclosures through a tube tunnel system connected directly to their home enclosures. Both experimental enclosures were identically equipped with three sitting boards and several branches suspended from the top for the animals to walk on. On the outer side of each experimental enclosure, we set up a small metal table with two wooden boxes on top that could be opened and closed from the outside. These boxes contained two different types of stimuli: an ambiguous object (a furry toy animal with large eyes, Figure 7B) and food items (mealworms). We selected a toy as an ambiguous object because of its uncertain valence – while it is visually salient, the animals cannot immediately categorize it as benign or dangerous. Pilot observations of animals not participating in the study confirmed that they responded variably to the object: some approached and touched the stimulus without vocalizing, while others produced alertness calls when the toy was revealed, consistent with its ambiguous nature. These differences in response were what we sought to elicit, as they allowed us to investigate whether and how individuals signal their perceptions to a naïve partner who cannot see the object.

**Figure 7:**
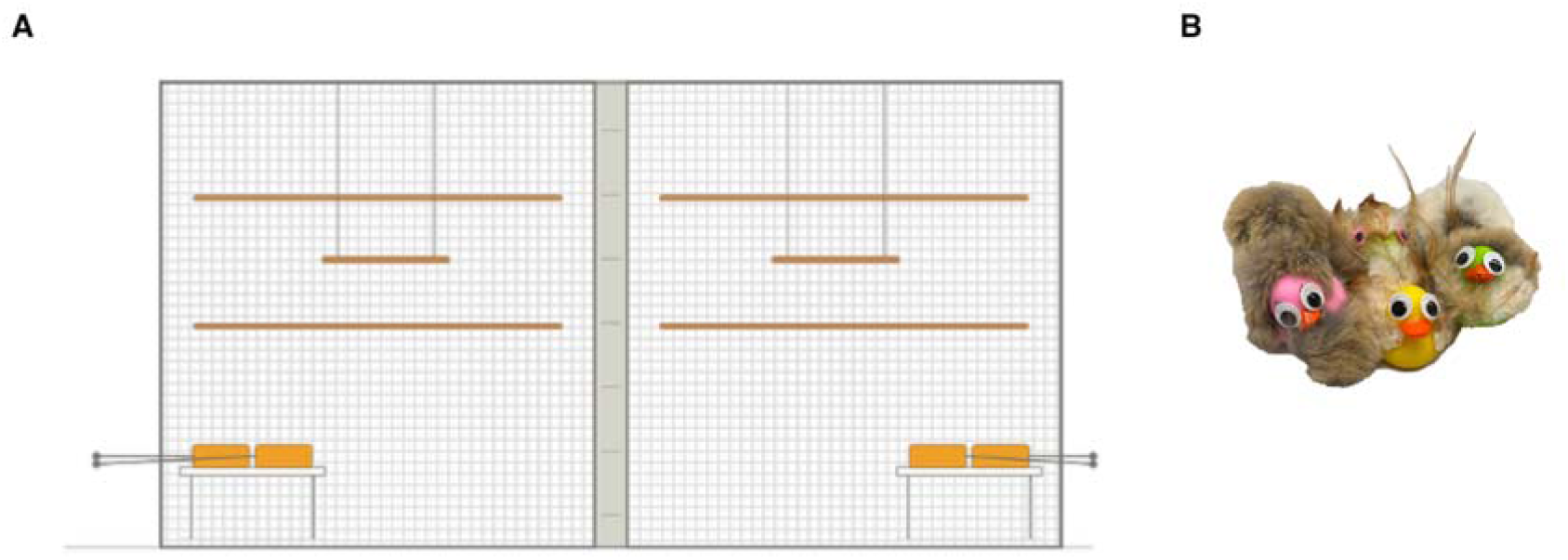
Experimental setup and ambiguous objects. **(A)** Experimental enclosures with access directly from home enclosures via a tube system. Metal tables with wooden boxes containing the ambiguous and food items are located at the outer end of each enclosure. Each enclosure is equipped with identical sitting boards and branches. An opaque screen can be placed in the gap separating the two enclosures to create visual isolation. **(B)** Examples of the ambiguous objects used in this study – differently colored rubber animal toys augmented with googly eyes, fur, or feathers, presented as a novel object of unclear identity and valence.

### Experimental procedure

We tested all possible dyads within each group. Prior to the experiments, all individuals were familiarized and habituated to the experimental enclosure by spending 2–3 sessions of approximately 20 minutes in it as a group, with some food provided on the tables. During the experiments, each individual was exposed to six different test conditions. Each session began with dyad partners having acoustic and visual contact for one minute (visual condition) to ensure that each individual was aware of who their partner was prior to introducing information asymmetry. In the following condition, the experimenter placed an opaque screen between the two compartments, blocking visual contact (non-visual neutral condition). This screen remained in place for the rest of the session, and all subsequent conditions occurred in visual (but not acoustic) occlusion (non-visual conditions). The neutral visual separation lasted for five minutes. The following four conditions, each lasting three minutes, were presented in a randomized order for each test session and dyad to reduce potential order and habituation effects. In each of these conditions, the experimenter opened one of the wooden boxes, which contained either an ambiguous item or food, for one of the subjects. The subject whose box was opened could approach and interact with its contents freely. Their partner in the other compartment remained passive, able to hear only any eventual vocalizations from the individual interacting with the items in the box. After three minutes of unrestricted access to the box’s contents, the experimenter closed the current box and opened the next one. The same procedure was repeated for the remaining conditions, ensuring that both individuals interacted with the ambiguous stimuli and the food. An overview of the experimental conditions is shown in Table 1. Each dyad was tested twice (pairs without offspring three times), resulting in a total of 119 test sessions.

**Table 1:**
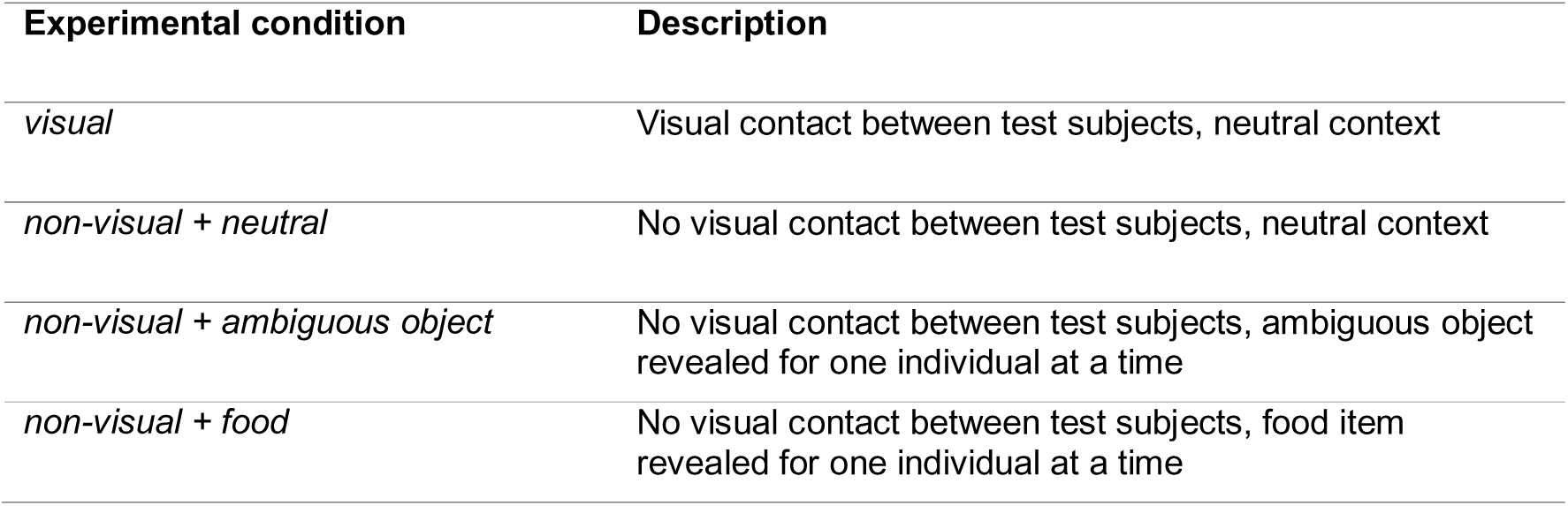
Experimental conditions during test sessions.

| Experimental condition | Description |
| --- | --- |
| *visual* | Visual contact between test subjects, neutral context |
| *non-visual + neutral* | No visual contact between test subjects, neutral context |
| *non-visual + ambiguous object* | No visual contact between test subjects, ambiguous object revealed for one individual at a time |
| *non-visual + food* | No visual contact between test subjects, food item revealed for one individual at a time |

### Acoustic recording

We recorded all vocalizations produced by test subjects throughout the entire test session using two AviSoft Bioacoustics CM16/CMPA microphones, each connected to an AviSoft Ultrasound Gate 116 at a sampling rate of 192 kHz. The microphones were set to different gain settings to ensure that both very loud (low gain) and quieter (high gain) vocalizations were captured with good acoustic quality. Both microphones were positioned on a tripod 1.5 meters from the experimental enclosures. All vocalizations were annotated in real time by assigning keyboard keys to the focal individuals (“left”/”right” according to the side of the enclosure where each vocalizing individual was located) and pressing them whenever a subject vocalized, to create a timestamp with the caller’s ID.

### Further processing of the acoustic data

All annotated vocalizations were further processed using AviSoft-SAS-Lab Pro (Version 5.3.01). We defined a call as a single acoustic element, meaning we broke down bouts and compound calls into individual syllables. We manually assigned call types based on their time-frequency representations^21,23,28^ and the caller’s identity as determined by the initial timestamp annotation for each element. 10% of all recordings were coded by two raters to assess inter-rater reliability (K = 0.846, n = 5,733, see Supplementary Methods S1 & Supplementary Figure S1, Supplementary Results S1), with agreement varying by call type (rarer call types showing lower agreement. We collected a total of 69,538 vocalizations throughout all sessions, which we grouped into three broader functional categories^21,22^ (Table 2). We based the grouping on our a priori predictions about which call types would show increased production rates under the specific experimental conditions. Contact-seeking vocalizations included “phee*”* (>0.5s), “short phees” (0.3–0.5s), “phee-peeps” (<0.3s), “trills”, “trill-peeps” (<0.2s), “trill-phees*”*, and “twitters*”*. Alertness calls included “mobbing calls” (“tsik*”*, “ekk*”*, “ock*”*) and “alarm calls*”*. Finally, food-indicating calls comprised “food chirps*”*. Spectrograms of some examples are shown in Figure 8. Call types that could not be assigned based on the definitions we used were put into a category “other” (less than 1%, Table 2) and only used when analyzing total volubility over all calls lumped together. Analyzing the calls at the functional category level was a statistical necessity, as several call types occurred at rates too low to permit reliable individual-level modeling.

**Table 2:** Distribution of recorded call types across call type categories. Total number of calls recorded per call type across all experimental conditions, sessions, and individuals (N = 69,538 total calls), with the proportion of the overall call sample each type represents.

| Call type | Category | N | Proportion |
| --- | --- | --- | --- |
| phee | contact | 10952 | 15.7% |
| short phee | contact | 1059 | 1.5% |
| phee-peep | contact | 2438 | 3.5% |
| trill | contact | 396 | 0.6% |
| trill-phee | contact | 363 | 0.5% |
| trill-peep | contact | 275 | 0.4% |
| twitter | contact | 2249 | 3.2% |
| tsik | alertness | 11615 | 16.7% |
| ekk | alertness | 29757 | 42.8% |
| ock | alertness | 1641 | 2.4% |
| alarm call | alertness | 63 | 0.1% |
| food chirp | food | 8131 | 11.7% |
| other | NA | 599 | 0.9% |

**Figure 8:**
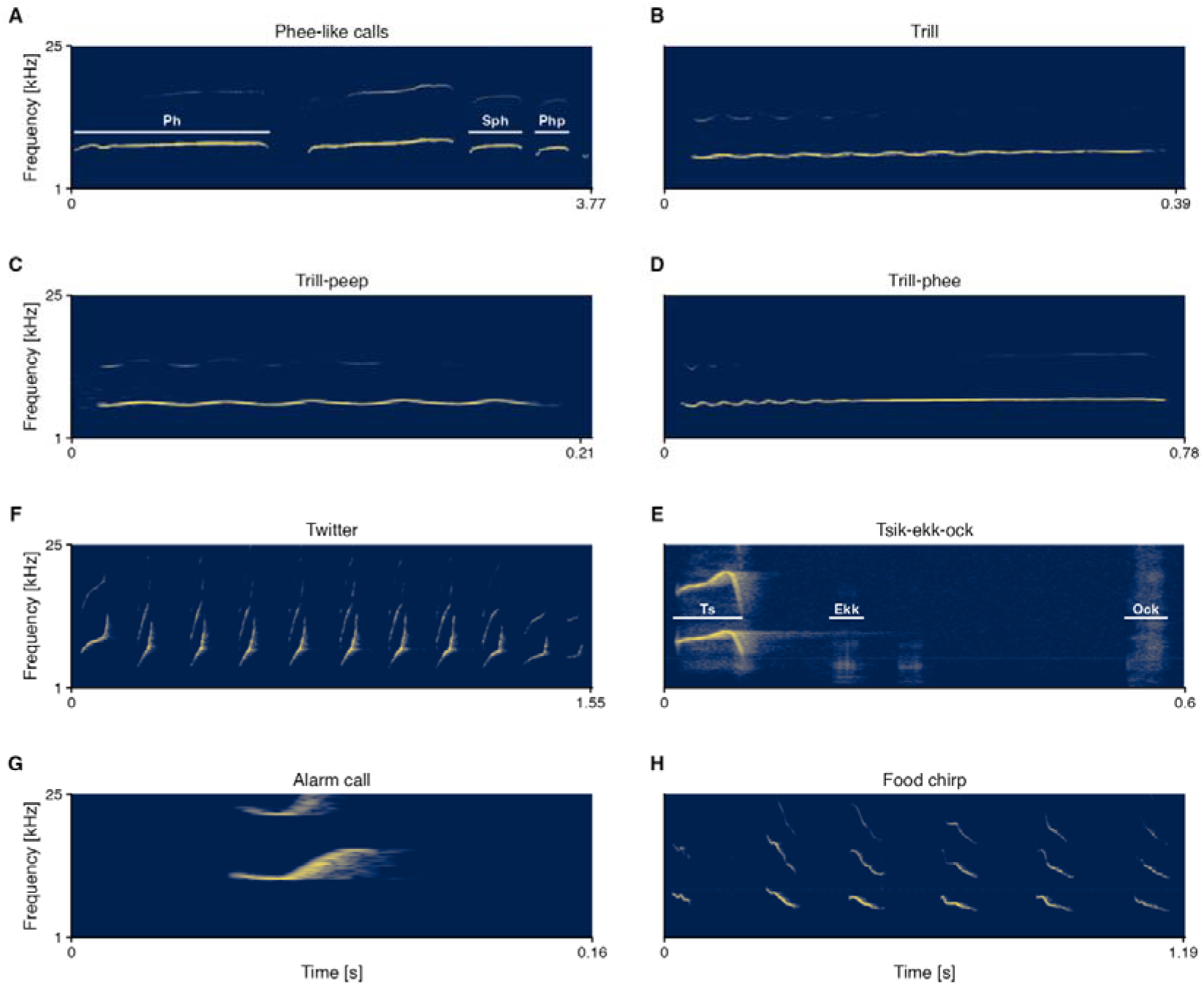
Spectrograms of common marmoset call types. Representative examples of the call types recorded in this study: **(A)** phee-like calls, comprising phee (Ph), short phee (Sph), and phee-peep (Php) syllables; **(B)** trill; **(C)** trill-peep; **(D)** trill-phee; **(E)** tsik-ekk-ock, comprising tsik (Ts), ekk (Ekk), and ock (Ock) syllables; **(F)** twitter; **(G)** alarm call; and **(H)** food chirps. All spectrograms were computed using a Hanning window. Color intensity reflects spectral amplitude (darker = lower, brighter = higher energy).

### Statistical analyses

All statistical analyses were performed in RStudio (Version 2026.1.0.392^56^) using generalized linear mixed-effect models (GLMMs) with the *glmmTMB* package^57^. Calling rates were analyzed as overdispersed count data using a negative binomial error distribution. Because several call-type categories had excess zero observations, most models included a zero-inflation component. Cases of overdispersion were adjusted by adding an overdispersion component. To test specific hypotheses about differences in calling rates across call type categories and experimental conditions, we applied a priori-defined custom contrasts (see Supplementary Table S2). For models 1, 2, and 3A, they test whether call rates differ between the visual and non-visual conditions, between conditions where the caller directly interacts with a stimulus versus conditions in which their partner does, and between the two active stimulus conditions; for model 3A, they additionally test whether a subjects directly approaching of the ambiguous object increases their alertness call rate. For model 4A, the contrasts test whether food call rates differ between the food-for-self and food-for-partner conditions and whether the manner of interaction with the food predicts call rates. Model assumptions and fit were evaluated using simulation-based diagnostic tests from the *DHARMa* package^58^, including tests for uniformity, dispersion, outliers, and zero-inflation (see Supplementary Table S3).

For each analysis, we first fitted a biologically relevant full model comprising two analytical layers. A confirmatory layer included the a priori condition contrasts (Supplementary Table S2), and individual predictors (sex, status) specified a priori based on our hypotheses; these were retained in all models regardless of significance. An exploratory layer comprised two-and three-way interactions between caller sex, status, and condition to test whether specific individual characteristics modulated condition effects (where applicable). Because we made no directional predictions for these interactions, we pruned non-significant ones to obtain parsimonious additive models. Where interactions were significant, we additionally compared the interaction model’s fit to a purely additive model using likelihood-ratio tests and AIC values and retained interaction terms only when the interaction model provided a significantly better fit than the additive model. The pruning was thus restricted to the exploratory layer and did not affect the confirmatory contrasts. All models included random intercepts for individuals nested within family groups to account for repeated measures and controlled for age (in months) and test session number to account for developmental and habituation effects. Because experimental conditions differed slightly in duration, an offset term was included to account for variation in sampling effort. We compared the full, reduced, and null models using likelihood-ratio tests and AIC values (model-comparison results in Supplementary Table S4; full unpruned model outputs are provided in Supplementary Table S5).

We assessed how marmosets’ total volubility varied across conditions and individual characteristics by fitting a model with the total number of calls per test session, combining all type categories, as the response variable (model 1). Fixed effects included caller sex, status, experimental condition (visual, non-visual neutral, ambiguous-for-self/partner, food-for-self/partner), caller age, and session number. To examine condition-specific differences in volubility by call type category, we fitted a separate model for contact calls (model 2) using the same predictor structure and random effects as the total volubility model. Furthermore, to analyze alertness call production, we constructed a condition variable that retains the six experimental conditions but subdivides the ambiguous-object-for-self condition into two levels, depending on whether the caller approached the stimulus or not (ambiguous-for-self with approach and ambiguous-for-self without approach). This allowed us to simultaneously test condition-level variation in alertness call rates as well as the effect of direct stimulus interaction, and to apply planned contrasts across all levels within a single model (model 3A, Supplementary Table S2). Fixed effects included caller sex, status, experimental condition, caller age, and session number. After pruning non-significant interactions from the exploratory level, the model retained a significant interaction between caller status and experimental condition. We were furthermore interested in how a subject’s alertness calls in the ambiguous-for-self condition might affect their naive partner’s vocal responses. To address this, we fitted a model with partner alertness calls as the response variable, adding the subject’s alertness calls, sex, status, and age as fixed effects, while controlling for test session number (model 3B, contagion hypothesis). Because partners’ responses leveled off at higher subject call rates, we log-transformed the subject’s call count. This greatly improved model fit compared with using untransformed counts (ΔAIC = 16.2). Additionally, we examined whether subject alertness calls predicted partner contact calls (de-escalation hypothesis), using the same predictor structure with partner contact calls as the response variable (model 3C).

To examine food call production, we restricted the dataset to food conditions only (food-for-self and food-for-partner), and constructed a four-level condition variable distinguishing food-for-self with feeding (subject approached the food and fed on it), food-for-self without feeding (subject only approached the food), food-for-self without approach, and food-for-partner, allowing us to apply planned contrasts across all levels within a single model (model 4A). We fitted a model with food calls as the response variable, adding caller sex, status, age, and session number as fixed effects. As with alertness calls, we were interested in what effect a subject’s food calls might have on the vocal behavior of their partner, specifically whether food calls could predict contact calling in the naïve partner (model 4B) and whether they would trigger increased alertness calls as a manifestation of inequity aversion in the partner without food access (model 4C). Both models included the subject’s food calls, sex, status, age, and session number as fixed effects, with model 4B additionally including an interaction between subject food calls and subject status that remained significant after pruning the exploratory layer; partner contact calls and partner alertness calls served as the respective response variables.

To address the potential confound arising from the fixed order of the visual and non-visual conditions, we additionally conducted a supplementary trend analysis adding condition order to models 1, 2, and 3A; model outputs are provided in Supplementary Table S6, Supplementary Methods S2, and Supplementary Results S2.

We chose call rate as our primary measure of volubility because it directly reflects the number of communicative acts produced per unit time, which we consider the most appropriate operationalization of volubility as a propensity to vocalize frequently; robustness checks comparing call rate to proportion of time spent vocalizing are provided in the Supplementary Materials (Supplementary Figure S2, Supplementary Tables S7 and S8, Supplementary Methods S3 and Supplementary Results S3).

## Declarations

### Ethics approval and consent to participate

The experiments were approved by the Kantonales Veterinäramt Zürich, Switzerland (license number ZH 232/19), and comply with Swiss law.

### Consent for publication

Not applicable

### Availability of data and materials

The datasets generated and/or analyzed during the current study are available on OSF, https://osf.io/7dsxh.

### Authors’ contributions

M.M.: Conceptualization, Methodology, Investigation, Data curation, Formal analysis, Visualization, Project administration, Writing – original draft, Writing – review & editing. R.K.B.: Data curation, Formal analysis, Visualization, Writing – review & editing. J.M.B.: Conceptualization, Methodology, Supervision, Resources, Funding acquisition, Writing – review & editing.

## Acknowledgements

This work has received funding from the European Research Council (ERC) under the European Horizon 2020 research and innovation program (Grant agreement No. 101001295).

We thank Alyssa Müller for her valuable assistance in annotating the vocal data. Furthermore, we are grateful to the animal keepers, Hidir Sengül and Dominique Ziegler, for their instrumental role in keeping the animals healthy and well cared for.

