## Supplementary Materials for "Volubility and vocal coordination under information asymmetry in captive common marmosets (*Callithrix jacchus*)"

Table S1: Study subjects participating in the study.

Table S2: A priori planned contrasts for GLMMs 1, 2, 3A, and 4A.

Table S3: Simulation-based model diagnostic results for fitted GLMMs.

Table S4: Model comparison statistics for fitted GLMMs.

Table S5: Type II analysis of deviance for all full unpruned and reduced GLMMs, showing robustness of reported effects to inclusion of interaction terms.

Table S6: Type II Analysis of Deviance for supplementary condition order trend analyses.

Table S7: Type II Analysis of Deviance for full and reduced GLMMs (Time spent vocalizing).

Table S8: Coefficient estimates from reduced models for proportion of time spent vocalizing.

Figure S1: Confusion matrix heatmap for inter-rater-reliability (row-wise normalized).

Figure S2: Relationship between call rate and proportion of time spent vocalizing across call-type categories.

Figure S3: Habituation effect on call rate across test sessions.

Supplementary Methods S1: Inter-rater reliability

Supplementary Methods S2: Condition order trend analysis

Supplementary Methods S3: Time spent vocalizing

Supplementary Results S1: Inter-rater reliability

Supplementary Results S2: Condition order trend analysis

Supplementary Results S3: Call rate vs. proportion of time spent vocalizing

**Table S1: Study subjects participating in the study.** List of animals by group, along with information about sex, status, and date of birth. f = female, m = male, b = breeder, h = helper

| Group | Individual | Sex | Status | Birth Date |
| --- | --- | --- | --- | --- |
| Gingers | Ginger  Jam | f  m | b  b | 04.05.2018  01.07.2016 |
| Grappas | Grappa  Craken  Gina  Giotto | f  m  f  m | b  b  h  h | 17.03.2015  10.10.2013  11.11.2019  11.11.2019 |
| Guapas | Guapa  Ninja  Guarana  Gimli | f  m  f  m | b  b  h  h | 04.05.2018  12.04.2015  05.03.2021  05.03.2021 |
| Jajas | Jaja  Membo  Jala  Jelly | f  m  f  m | b  b  h  h | 29.08.2009  17.06.2009  27.01.2015  27.01.2015 |
| Jambis | Jambi  Werewolf  Jafar  Jaggar | f  m  m  m | b  b  h  h | 10.11.2015  10.05.2018  07.07.2021  07.07.2021 |
| Limas | Lima  Wolverine  Lychee  Lemon | f  m  f  m | b  b  h  h | 15.02.2015  10.05.2018  20.10.2021  20.10.2021 |
| Mibbas | Mibba  Conan  Madame  Madita | f  b  f  f | b  b  h  h | 17.06.2009  10.10.2013  08.04.2018  08.04.2018 |
| Mojitas | Mojita  Umberto | f  m | b  b | 04.05.2015  2016/2017 |
| Mulans | Mulan  Odin | f  m | b  b | 26.05.2017  25.02.2016 |
| Nocciolettas | Noccioletta  Ulysses | f  m | b  b | 18.11.2019  17.08.2019 |
| Olympias | Olympia  Wuschel | f  m | b  b | 25.02.2016  23.10.2019 |
| Wasabis | Wasabi  Lynx  Wakame  Wonton | f  m  f  f | b  b  h  h | 14.01.2017  23.11.2018  03.10.2021  03.10.2021 |
| Washingtons | Washington  Lotus  Wall E  Woody | f  m  f  m | b  b  h  h | 30.08.2013  03.07.2012  12.08.2021  12.08.2021 |

**Table S2: A priori planned contrasts for GLMMs 1, 2, 3A, and 4A.** List of planned contrasts and their conceptual definitions used to test specific hypotheses.

| Models 1 & 2: All calls and contact-seeking calls (all conditions, 6 levels) | |
| --- | --- |
| *visual vs. remaining* | Visual condition against all conditions of visual separation regardless of whether an individual gets to interact with a stimulus or not  *Test effect of visual access between dyadic partners on calling in general and contact calls specifically* |
| *non-visual neutral vs. stimulus conditions* | Neutral non-visual condition against non-visual conditions where an individual could interact with a stimulus  *Test effect of stimulus presence vs. absence under visual separation between dyadic partners on calling in general and contact calls specifically* |
| *stimulus-for-self vs. stimulus-for-partner* | Non-visual conditions in which a subject interacts with a stimulus against conditions in which their partner interacts with a stimulus  *Test effect of information asymmetry between dyadic partners in conditions in which a stimulus is present on calling in general and contact calls specifically* |
| *food-for-self vs. ambiguous-object-for-self* | Non-visual condition in which a subject interacts with food against non-visual condition in which the subject interacts with an ambiguous stimulus  *Test effect of stimulus type when having access to the stimulus on calling in general and contact calls specifically* |
| *food-for-partner vs. ambiguous-object-for-partner* | Non-visual condition in which a subject’s partner interacts with food against non-visual condition in which the partner interacts with an ambiguous stimulus  *Test effect of stimulus type when partner has access to the stimulus on calling in general and contact calls specifically* |

**Model 3A: Alertness calls (all conditions, 7 levels)**

| *visual vs. remaining* | Visual condition against all conditions of visual separation regardless of whether an individual gets to interact with a stimulus or not.  *Test effect of visual access between dyadic partners on alertness calling* |
| --- | --- |
| *non-visual neutral vs. stimulus conditions* | Neutral non-visual condition against non-visual conditions where an individual could interact with a stimulus  *Test effect of stimulus presence vs. absence under visual separation between dyadic partners on alertness calling* |
| *ambiguous object vs. food* | Non-visual ambiguous object conditions (pooled across self/partner) against non-visual food conditions (pooled across self/partner)  *Test effect of stimulus type on alertness calling* |
| *ambiguous-object-for-self vs. ambiguous-object- for-partner* | Non- visual ambiguous object condition in which a subject may interact with the object (pooled across approach categories, i.e., approach object vs. no approach) against non-visual ambiguous object condition in which their partner may interact with the object.  *Test effect of information asymmetry in ambiguous stimulus conditions on alertness calling* |
| *food-for-self vs. food-for-partner* | Non-visual condition in which a subject may interact with food (mealworms) against a non-visual condition in which their partner may interact with food.  *Test effect of information asymmetry in food conditions (null expectation for alertness calls)* |
| *approach vs. no approach* | Within the ambiguous object for self-condition, subjects who approached the ambiguous stimulus against those who did not.  *Test effect of direct interaction with the ambiguous object on alertness calling* |

**Model 4A: Food calls (food conditions, 4 levels)**

| *food-for-self vs. food-for-partner* | Food-for-self conditions (pooled across interaction with food categories, i.e., feed, only approach, no approach) against the food for partner conditions  *Test effect of information asymmetry between dyadic partners in food conditions of food calling* |
| --- | --- |
| *approach and feed vs. remaining* | Within the food-for-self condition, subjects who approached and fed on the food against cases when they only approached or did not approach at all  *Test effect of food consumption on food calling* |
| *only approach vs. no approach* | Within the food-for-self condition, subjects who approached the food but did not consume any of it against cases when they did not approach at all  *Test effect of awareness that food is present without consumption on food calling* |

**Table S3: Simulation-based model diagnostic results for fitted GLMMs.** DHARMa residual diagnostics (n = 1000 simulations) for each full and reduced model, including the Kolmogorov-Smirnov (KS) test for residual uniformity, dispersion, zero-inflation, and outlier tests (all based on simulated residuals). Significant p-values are shown in bold. Statistical trends are indicated by a point (•).

|  | **KS-test** | **Dispersion** | **Zero-inflation** | **Outlier** |
| --- | --- | --- | --- | --- |
| **Model 1: All calls across conditions** | | | | |
| full | 0.448 | 0.404 | 0.058• | **0.023*** |
| reduced | 0.245 | 0.312 | 0.064• | **0.023*** |
| **Model 2: Contact calls across conditions** | | | | |
| full | 0.248 | 0.952 | 0.110 | 0.061• |
| reduced | 0.183 | 0.960 | 0.164 | 0.362 |
| **Model 3A: Alertness calls across conditions** | | | | |
| full | 0.677 | 0.202 | 0.198 | 0.757 |
| reduced | 0.742 | 0.424 | 0.162 | 0.060• |
| **Model 3B: Alertness contagion** | | | | |
| full | 0.382 | 0.830 | 1.000 | 1.000 |
| reduced | 0.737 | 0.872 | 0.952 | 1.000 |
| **Model 3C: Alertness calls subject and contact calls partner** | | | | |
| full | 0.647 | 0.774 | 0.258 | 1.000 |
| reduced | 0.487 | 0.638 | 0.996 | 1.000 |
| **Model 4A: Food calls across food conditions** | | | | |
| full | 0.544 | 0.402 | 0.574 | 1.000 |
| reduced | 0.864 | 0.296 | 0.444 | 1.000 |
| **Model 4B: Food calls subject and contact calls partner** | | | | |
| full | 0.465 | 0.638 | 0.334 | 1.000 |
| reduced | 0.766 | 0.714 | 0.354 | 1.000 |
| **Model 4C: Food calls subject and alertness calls partner** | | | | |
| full | 0.363 | 0.960 | 0.532 | 1.000 |
| reduced | 0.531 | 0.994 | 0.550 | 1.000 |

**Table S4: Model comparison statistics for fitted GLMMs.** Likelihood ratio test comparisons among null, reduced, and full models, reporting AIC, BIC, $\chi$², degrees of freedom (Df), and p-values. Significant p-values are shown in bold. Statistical trends are indicated by a point (•).

|  | **AIC** | **BIC** | $\boldsymbol{\chi}$**²** | **Df** | ***p*-value** |
| --- | --- | --- | --- | --- | --- |
| **Model 1: All calls across conditions** | | | | | |
| null | 12776.6 | 12797 | - | - | - |
| reduced | 12382.8 | 12492.6 | 427.839 | 17 | **<.001 ***** |
| full | 12393.1 | 12586.6 | 21.614 | 16 | 0.156 |
| **Model 2: Contact calls across conditions** | | | | | |
| null | 9485.3 | 9506.2 | - | - | - |
| reduced | 9165.5 | 9270.1 | 351.855 | 16 | **<.001 ***** |
| full | 9182.9 | 9371.2 | 14.525 | 16 | 0.560 |
| **Model 3A: Alertness calls across conditions** | | | | | |
| null | 8807.1 | 8828.0 | - | - | - |
| reduced | 8611.1 | 8752.6 | 241.463 | 23 | **<.001 ***** |
| full | 8620.7 | 8829.6 | 16.946 | 13 | 0.202 |
| **Model 3B: Alertness contagion** | | | | | |
| null | 1392.9 | 1406.5 | - | - | - |
| reduced | 1346.4 | 1383.8 | 60.491 | 7 | **<.001 ***** |
| full | 1353.8 | 1404.8 | 0.638 | 4 | 0.959 |
| **Model 3C: Alertness calls subject and contact calls partner** | | | | | |
| null | 1557.9 | 1571.5 | - | - | - |
| reduced | 1548.0 | 1592.2 | 27.888 | 9 | **0.004**** |
| full | 1546.7 | 1597.7 | 5.266 | 2 | 0.072• |
| **Model 4A: Food calls across food conditions** | | | | | |
| null | 1655.5 | 1671.8 | - | - | - |
| reduced | 1464.2 | 1517.4 | 209.252 | 9 | **<.001 ***** |
| full | 1466.0 | 1523.3 | 0.206 | 1 | 0.650 |
| **Model 4B: Food calls subject and contact calls partner** | | | | | |
| null | 1535.6 | 1549.2 | - | - | - |
| reduced | 1504.8 | 1549.0 | 48.748 | 9 | **<.001 ***** |
| full | 1508.9 | 1563.2 | 1.978 | 3 | 0.577 |
| **Model 4C: Food calls subject and alertness calls partner** | | | | | |
| null | 1466.2 | 1479.8 | - | - | - |
| reduced | 1471.9 | 1509.3 | 8.339 | 7 | 0.304 |
| full | 1477.9 | 1528.8 | 2.029 | 4 | 0.730 |

**Table S5: Type II analysis of deviance for all full unpruned and reduced GLMMs**, **showing robustness of reported effects to inclusion of interaction terms.** Comparison of fixed effects and interaction terms across full and reduced models for fitted GLMMs reporting $\chi$², degrees of freedom (Df), and p-value. Significant p-values are shown in bold. Statistical trends are indicated by a point (•).

|  | Full model | | | Reduced model | | |
| --- | --- | --- | --- | --- | --- | --- |
|  | $\boldsymbol{\chi}$**²** | **Df** | ***p*-value** | $\boldsymbol{\chi}$**²** | **Df** | ***p*-value** |
| Model 1: All calls across conditions | | | | | | |
| *sex* | 0.579 | 1 | 0.447 | 0.606 | 1 | 0.436 |
| *status* | 0.399 | 1 | 0.528 | 0.465 | 1 | 0.495 |
| *condition* | 97.675 | 5 | **<.001***** | 96.038 | 5 | **<.001***** |
| *age (months* | 2.952 | 2 | 0.086• | 2.701 | 1 | 0.100 |
| *session* | 13.020 | 1 | **<.001***** | 13.174 | 1 | **<.001***** |
| *sex : status* | 0.002 | 1 | 0.966 | - | - | - |
| *sex : condition* | 7.663 | 5 | 0.176 | - | - | - |
| *status : condition* | 10.416 | 5 | 0.064• | - | - | - |
| *sex : status : condition* | 5.014 | 5 | 0.414 | - | - | - |
| Model 2: Contact calls across conditions | | | | | | |
| *sex* | 1.139 | 1 | 0.286 | 1.122 | 1 | 0.289 |
| *status* | 3.532 | 1 | 0.060• | 3.592 | 1 | 0.058• |
| *condition* | 4.923 | 5 | 0.425 | 5.172 | 5 | 0.395 |
| *age (months)* | 0.735 | 1 | 0.391 | 0.521 | 1 | 0.470 |
| *session* | 8.243 | 1 | **0.004**** | 8.145 | 1 | **0.004**** |
| *sex : status* | 0.329 | 1 | 0.566 | - | - | - |
| *sex : condition* | 5.081 | 5 | 0.406 | - | - | - |
| *status : condition* | 6.594 | 5 | 0.253 | - | - | - |
| *sex : status : condition* | 2.108 | 5 | 0.834 | - | - | - |
| Model 3A: Alertness calls across conditions | | | | | | |
| *sex* | 0.317 | 1 | 0.574 | 0.332 | 1 | 0.565 |
| *status* | 1.026 | 1 | 0.311 | 0.958 | 1 | 0.328 |
| *condition* | 21.830 | 6 | **0.001**** | 19.386 | 6 | **0.004**** |
| *age (months)* | 5.192 | 1 | **0.023*** | 5.220 | 1 | **0.022*** |
| *session* | 7.424 | 1 | **0.006**** | 5.887 | 1 | **0.015*** |
| *sex : status* | 1.142 | 1 | 0.285 | - | - | - |
| *sex : condition* | 9.472 | 6 | 0.149 | - | - | - |
| *status : condition* | 35.682 | 6 | **< .001***** | 32.853 | 6 | **< .001***** |
| *sex : status : condition* | 6.093 | 6 | 0.413 | - | - | - |
| Model 3B: Alertness contagion | | | | | | |
| *alertness calls (log)* | 39.561 | 1 | **< .001***** | 41.035 | 1 | **< .001***** |
| *sex* | 0.060 | 1 | 0.806 | 0.049 | 1 | 0.826 |
| *status* | 0.101 | 1 | 0.751 | 0.095 | 1 | 0.757 |
| *age (months)* | 0.494 | 1 | 0.482 | 0.624 | 1 | 0.430 |
| *session* | 0.115 | 1 | 0.735 | 0266 | 1 | 0.606 |
| *alertness calls (log) : sex* | 0.165 | 1 | 0.685 | - | - | - |
| *alertness calls (log) : status* | 0.164 | 1 | 0.686 | - | - | - |
| *sex : status* | 0.327 | 1 | 0.568 | - | - | - |
| *alertness calls (log) : sex : status* | 0.068 | 1 | 0.794 | - | - | - |
| Model 3C: Alertness calls subject and contact calls partner | | | | | | |
| *alertness calls* | 1.476 | 1 | 0.224 | 1.050 | 1 | 0.306 |
| *sex* | 0.806 | 1 | 0.369 | 2.279 | 1 | 0.131 |
| *status* | 3.066 | 1 | 0.080• | 1.012 | 1 | 0.314 |
| *age (months)* | 0.083 | 1 | 0.774 | 0.203 | 1 | 0.652 |
| *session* | 10.026 | 1 | **0.002**** | 10.644 | 1 | **0.001**** |
| *alertness calls : sex* | 3.905 | 1 | **0.048*** | - | - | - |
| *alertness calls : status* | 0.267 | 1 | 0.605 | - | - | - |
| *sex : status* | 0.822 | 1 | 0.365 | - | - | - |
| *alertness calls : sex : status* | 0.006 | 1 | 0.939 | - | - | - |
| Model 4A: Food call across food conditions | | | | | | |
| *sex* | 3.009 | 1 | 0.083• | 5.198 | 1 | **0.023*** |
| *status* | 0.076 | 1 | 0.783 | 0.063 | 1 | 0.802 |
| *condition* | 129.707 | 3 | **< .001***** | 137.749 | 3 | **< .001***** |
| *age (months)* | 0.047 | 1 | 0.829 | 0.116 | 1 | 0.734 |
| *session* | 1.196 | 1 | 0.274 | 1.169 | 1 | 0.280 |
| *sex : status* | 0.181 | 1 | 0.670 | - | - | - |
| Model 4B: Food calls subject and contact calls partner | | | | | | |
| *food calls* | 1.631 | 1 | 0.202 | 2.140 | 1 | 0.144 |
| *sex* | 0.035 | 1 | 0.852 | 0.027 | 1 | 0.870 |
| *status* | 6.337 | 1 | **0.012*** | 4.929 | 1 | **0.026*** |
| *age (months)* | 0.027 | 1 | 0.870 | 0.064 | 1 | 0.800 |
| *session* | 2.412 | 1 | 0.120 | 0.064 | 1 | 0.800 |
| *food calls : sex* | 1.148 | 1 | 0.284 | - | - | - |
| *food calls : status* | 4.196 | 1 | **0.041*** | 9.628 | 1 | **0.002**** |
| *sex : status* | 0.449 | 1 | 0.503 | - | - | - |
| *food calls : sex : status* | 0.323 | 1 | 0.570 | - | - | - |
| Model 4C: Food calls subject and alertness calls partner | | | | | | |
| *food calls* | 0.030 | 1 | 0.862 | 0.000 | 1 | 0.993 |
| *sex* | 0.186 | 1 | 0.666 | 0.133 | 1 | 0.715 |
| *status* | 2.745 | 1 | 0.098• | 2.802 | 1 | 0.094• |
| *age (months)* | 4.323 | 1 | **0.038*** | 4.510 | 1 | **0.034*** |
| *session* | 0.060 | 1 | 0.806 | 0.016 | 1 | 0.900 |
| *food calls : sex* | 0.003 | 1 | 0.956 | - | - | - |
| *food calls : status* | 1.167 | 1 | 0.280 | - | - | - |
| *sex : status* | 0.651 | 1 | 0.420 | - | - | - |
| *food calls : sex : status* | 0.233 | 1 | 0.629 | - | - | - |

**Table S6: Type II Analysis of Deviance for supplementary condition order trend analyses.** Condition order was added as a polynomial factor to models 1, 2, and 3A, restricted to the four randomized conditions. Reporting $\chi$², degrees of freedom (Df), and p-values. Significant p-values are shown in bold. Statistical trends are indicated by a point (•).

|  | ***χ²*** | **Df** | ***p*-value** |
| --- | --- | --- | --- |
| **Model 1: All calls across randomized conditions (Trend analysis)** | | | |
| *sex* | 0.234 | 1 | 0.628 |
| *status* | 0.562 | 1 | 0.454 |
| *condition* | 76.332 | 3 | **< .001***** |
| *age (months)* | 2.112 | 1 | 0.146 |
| *session* | 19.874 | 1 | **< .001***** |
| *condition order* | 3.108 | 3 | 0.375 |
| **Model 2: Contact calls across randomized conditions (Trend analysis)** | | | |
| *sex* | 0.522 | 1 | 0.470 |
| *status* | 2.994 | 1 | 0.084• |
| *condition* | 2.681 | 3 | 0.443 |
| *age (months)* | 0.571 | 1 | 0.450 |
| *session* | 8.366 | 1 | **0.004**** |
| *condition order* | 3.818 | 3 | 0.282 |
| **Model 3A: Alertness calls across randomized conditions (Trend analysis)** | | | |
| *sex* | 0.241 | 1 | 0.623 |
| *status* | 0.736 | 1 | 0.391 |
| *condition* | 9.255 | 4 | 0.055• |
| *age (months)* | 5.462 | 1 | **0.019*** |
| *session* | 13.252 | 1 | **< .001***** |
| *condition order* | 8.810 | 3 | **0.032*** |
| *status : condition* | 32.270 | 4 | **< .001***** |

**Table S7: Type II Analysis of Deviance for full and reduced GLMMs (Time spent vocalizing).** Models 1 and 2 from the original analysis were replicated using the proportion of time spent vocalizing instead of call rate, modeled with the ordered-beta family. Reporting $\chi$², degrees of freedom (Df), and p-values. Significant p-values are shown in bold. Statistical trends are indicated by a point (•).

|  | Full model | | | Reduced model | | |
| --- | --- | --- | --- | --- | --- | --- |
|  | $\boldsymbol{\chi}$**²** | **Df** | ***p*-value** | $\boldsymbol{\chi}$**²** | **Df** | ***p*-value** |
| Model 1: All calls across conditions (Time spent vocalizing) | | | | | | |
| *sex* | 0.196 | 1 | 0.658 | 0.217 | 1 | 0.642 |
| *status* | 4.352 | 1 | **0.037*** | 4.423 | 1 | **0.035*** |
| *condition* | 53.123 | 5 | **< .001***** | 52.549 | 5 | **< .001***** |
| *age (months)* | 1.710 | 1 | 0.191 | 1.795 | 1 | 0.180 |
| *session* | 27.208 | 1 | **< .001***** | 27.000 | 1 | **< .001***** |
| *sex : status* | 0.152 | 1 | 0.697 | - | - | - |
| *sex : condition* | 9.994 | 5 | 0.075• | - | - | - |
| *status : condition* | 11.001 | 5 | 0.051• | - | - | - |
| *sex : status : condition* | 5.906 | 5 | 0.315 | - | - | - |
| Model 2: Contact calls across conditions (Time spent vocalizing) | | | | | | |
| *sex* | 1.139 | 1 | 0.286 | 1.122 | 1 | 0.289 |
| *status* | 3.532 | 1 | 0.060• | 3.592 | 1 | 0.058• |
| *condition* | 4.923 | 5 | 0.425 | 5.172 | 5 | 0.395 |
| *age (months)* | 0.735 | 1 | 0.391 | 0.521 | 1 | 0.470 |
| *session* | 8.243 | 1 | **0.004**** | 8.145 | 1 | **0.004**** |
| *sex : status* | 0.329 | 1 | 0.566 | - | - | - |
| *sex : condition* | 5.081 | 5 | 0.406 | - | - | - |
| *status : condition* | 6.594 | 5 | 0.253 | - | - | - |
| *sex : status : condition* | 2.108 | 5 | 0.834 | - | - | - |

**Table S8: Coefficient estimates from reduced models for proportion of time spent vocalizing.** Estimates, standard errors (SE), odds ratios (OR) with 95% confidence intervals (CI low and CI high), z-values, and p-values for all fixed effects in the reduced versions of models 1 and 2 using proportion of time spent vocalizing as the response variable. Significant p-values are shown in bold. Statistical trends are indicated by a point (•).

|  | **Estimate** | **SE** | **OR** | **CI low** | **CI high** | **z** | ***p*-value** |
| --- | --- | --- | --- | --- | --- | --- | --- |
| **Model 1: All calls across conditions** | | | | | | | |
| *male vs. female* | 0.059 | 0.127 | 1.061 | 0.827 | 1.361 | 0.466 | 0.642 |
| *helper vs. breeder* | 0.363 | 0.173 | 1.437 | 1.025 | 2.016 | 2.103 | **0.035*** |
| *visual vs. remaining conditions* | 0.031 | 0.012 | 1.031 | 1.008 | 1.055 | 2.624 | **0.009**** |
| *non-visual vs. stimuli conditions* | -0.023 | 0.010 | 0.977 | 0.958 | 0.996 | -2.348 | **0.019*** |
| *self vs. partner* | 0.097 | 0.024 | 1.102 | 1.053 | 1.154 | 4.144 | **< .001***** |
| *food for self vs. ambiguous for self* | 0.175 | 0.034 | 1.191 | 1.113 | 1.274 | 5.089 | **< .001***** |
| *food for other vs. ambiguous for other* | -0.011 | 0.032 | 0.989 | 0.929 | 1.053 | -0.348 | 0.728 |
| *age (months)* | -0.003 | 0.002 | 0.997 | 0.992 | 1.002 | -1.340 | 0.180 |
| session | -0.061 | 0.012 | 0.941 | 0.919 | 0.963 | -5.196 | **< .001***** |
| **Model 2: Contact calls across conditions** | | | | | | | |
| *male vs. female* | 0.178 | 0.168 | 1.195 | 0.860 | 1.660 | 1.059 | 0.289 |
| *helper vs breeder* | 0.452 | 0.238 | 1.571 | 0.985 | 2.507 | 1.895 | 0.058• |
| *visual vs remaining conditions* | -0.014 | 0.015 | 0.986 | 0.958 | 1.016 | -0.899 | 0.369 |
| *non-visual vs stimuli conditions* | -0.010 | 0.012 | 0.990 | 0.968 | 1.013 | -0.839 | 0.401 |
| *self vs partner* | -0.006 | 0.029 | 0.994 | 0.939 | 1.051 | -0.221 | 0.825 |
| *food for self vs ambiguous for self* | 0.075 | 0.040 | 1.078 | 0.997 | 1.165 | 1.896 | 0.058• |
| *food for other vs ambiguous for other* | 0.000 | 0.042 | 1.000 | 0.921 | 1.086 | 0.003 | 0.997 |
| *age (months)* | -0.003 | 0.004 | 0.998 | 0.991 | 1.004 | -0.722 | 0.470 |
| *session* | -0.046 | 0.016 | 0.955 | 0.925 | 0.986 | -2.854 | **0.004**** |

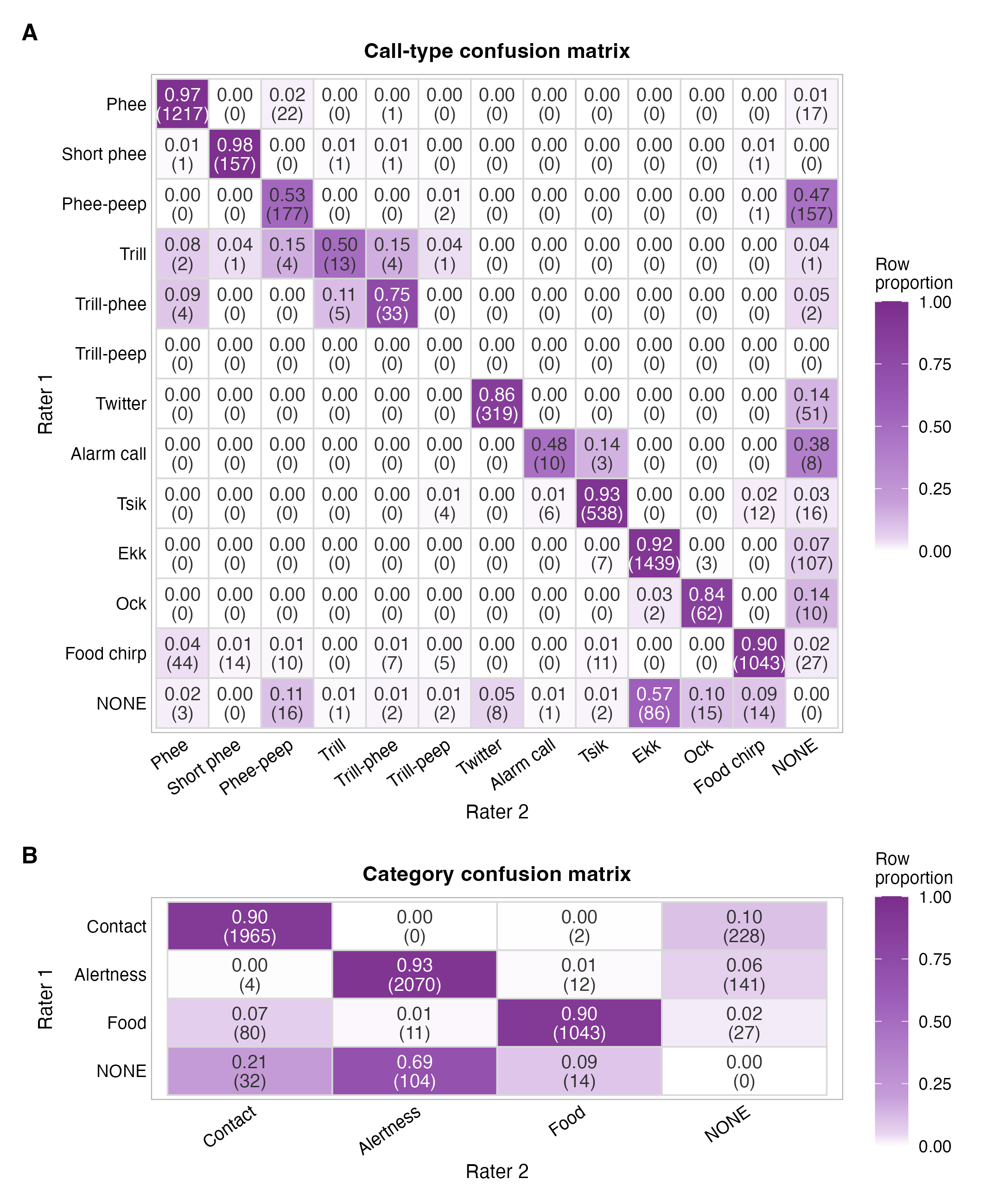

**Figure S1: Confusion matrix heatmap for inter-rater-reliability (row-wise normalized).** Heatmap showing agreement between the two raters for each annotated **(A)** call type and **(B)** functional call category. Cell values represent row-wise percentages (i.e., the proportion of Rater 1’s labels assigned to each label by Rater 2) and absolute call numbers in parentheses underneath. Darker shading indicates higher proportions.

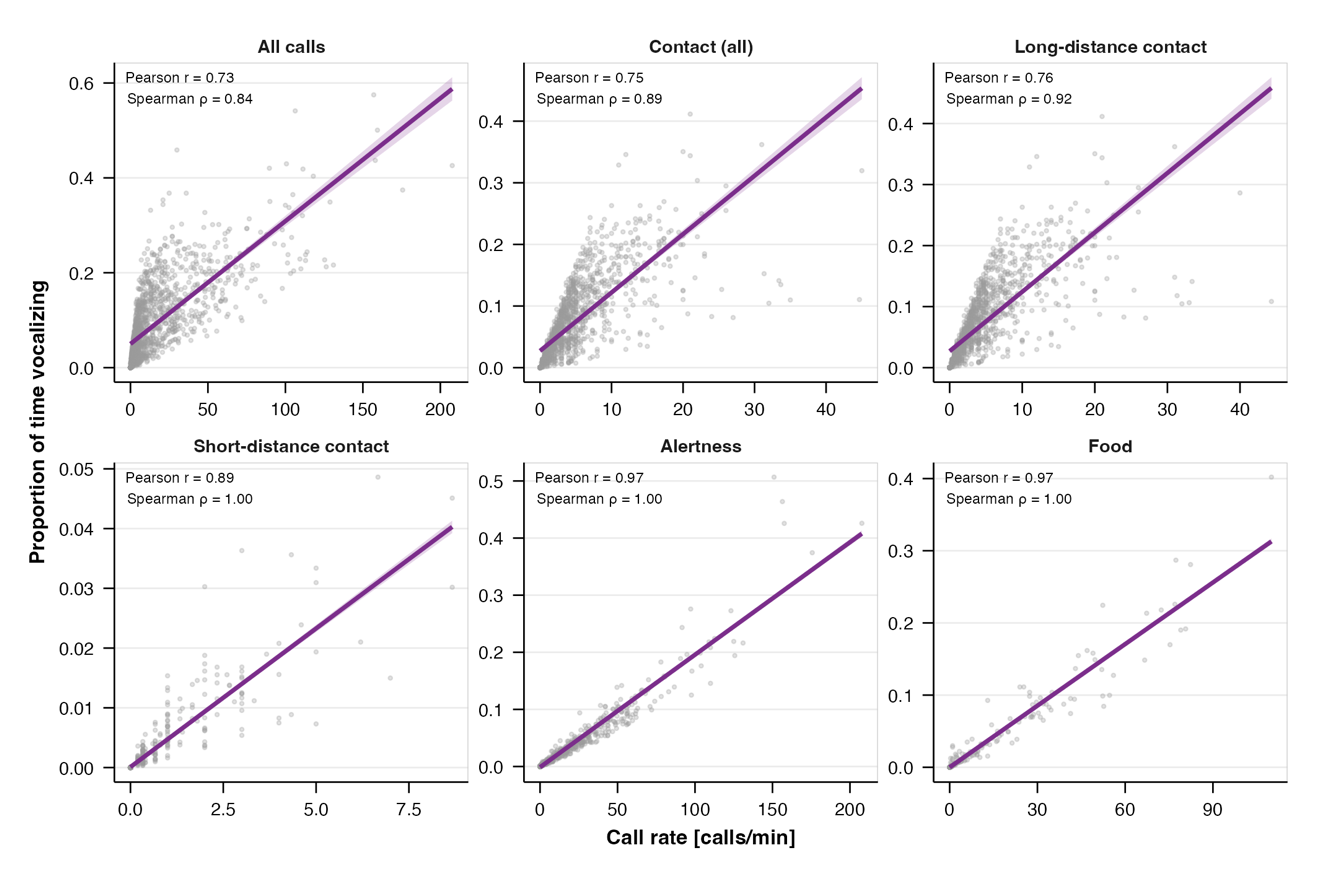

**Figure S2: Relationship between call rate and proportion of time spent vocalizing across call-type categories.** Each point represents one observation per caller, condition, and session (N = 1380 per panel). Purple lines and shaded areas show the linear fit and 95% confidence interval, respectively. Pearson and Spearman correlation coefficients are reported in each panel.

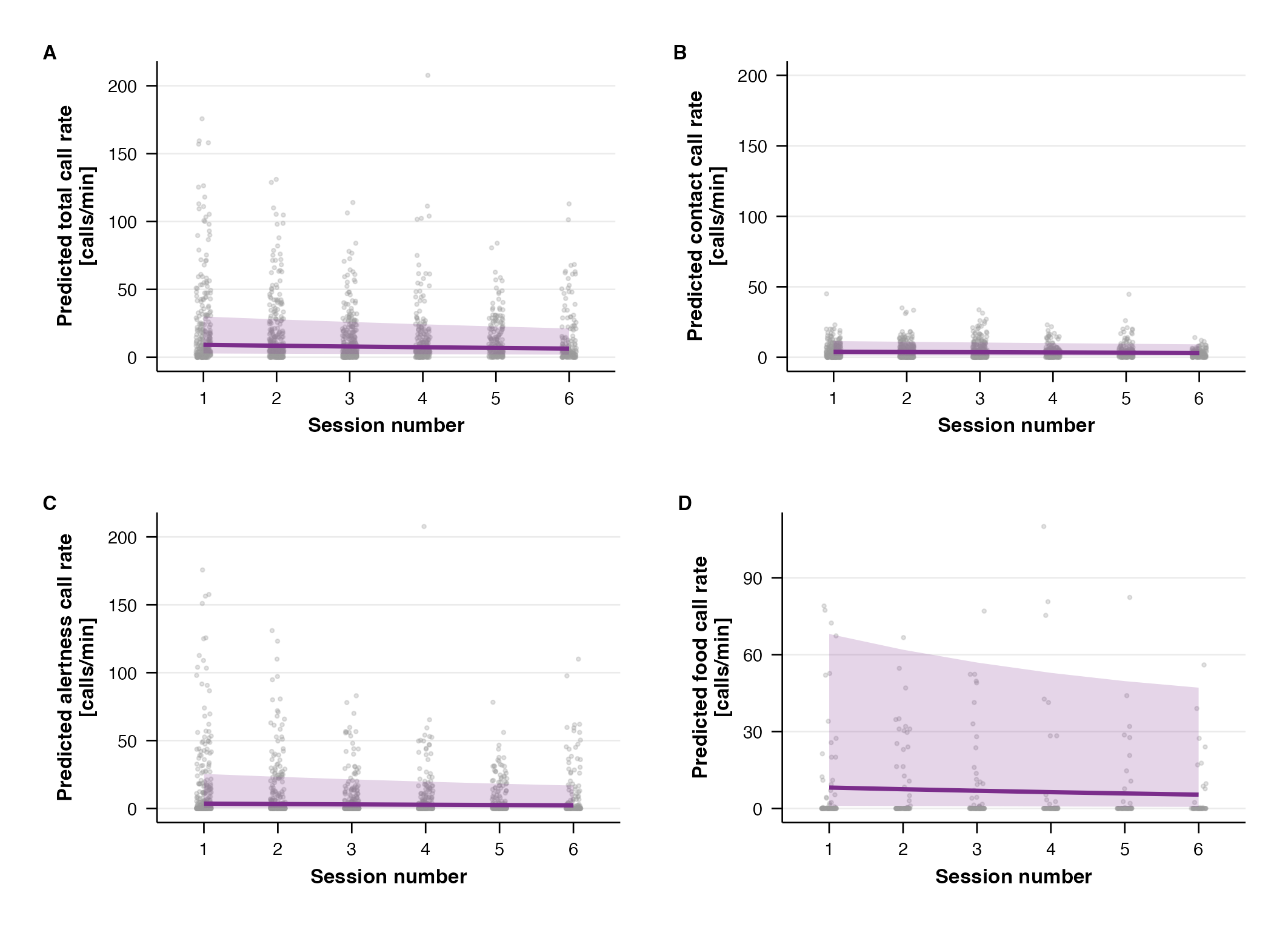

**Figure S3:** **Habituation effect on call rate across test sessions.** Model-predicted call rates as a function of session number for **(A)** all call types, **(B)** contact calls, **(C)** alertness calls, and **(D)** food calls. The purple lines and shaded areas indicate the fitted trend and 95% confidence interval, respectively. Dots represent individual observations.

Supplementary Methods S1: Inter-rater-reliability

We assessed inter-rater reliability using unweighted Cohen’s kappa. Segments were matched across raters using a temporal overlap criterion. For each recording, rater annotations were paired by maximizing Intersection over Union (IoU) of segment boundaries via the Hungarian algorithm ^1^, with a minimum IoU of 0.3 required for a valid match. Unmatched segments were retained as “none”-coded pairs, so kappa reflects agreement on both segmentation (whether a call was identified) and call-type classification.

Supplementary Methods S2: Condition order trend analysis

We addressed the potential confound arising from the fixed order of the visual and non-visual neutral conditions by conducting a supplementary trend analysis to test whether the within-session position of the experimental conditions influenced call rates. Because any linear decay in call rates due to habituation would be expected to carry over into subsequent conditions, we restricted the analysis to the four randomized conditions. We excluded the visual and non-visual neutral conditions, whose order was fixed by design, thereby precluding the separation of order from condition effects. We added condition order as a polynomial factor to models 1, 2, and 3A; model outputs are provided in Supplementary Table S6, and results are reported in Supplementary Results S2.

Supplementary Methods S3: Proportion of time spent vocalizing

Additionally, we examined whether our results were robust to the choice of vocal output measure. Liao et al. (2018)^2^ reported divergent patterns between call rate and time spent vocalizing across social-distance conditions in marmosets, raising the question of whether the two measures capture different aspects of vocal activity. We chose call rate as our primary measure of volubility because it directly reflects the number of communicative acts produced per unit time, which we consider the most appropriate operationalization of volubility as a propensity to vocalize frequently. However, divergence between the two measures is most likely to arise when call durations vary substantially, either across call types or within a call-type category, as longer calls would occupy more time per unit of time. We therefore focused this analysis on models 1 and 2, where such variation is most pronounced: model 1 combines all call types, which differ considerably in syllable duration, while model 2 covers contact calls that span from very long phee-syllables to short trills and phee-peeps. To assess whether variation in call duration caused the two measures to yield different results in our data, we replicated these models using the proportion of time spent vocalizing as the response variable, modeled with the ordered-beta family to accommodate the bounded-continuous nature of the proportion with a point mass at zero. The response variable, predictor structure, and random effects were identical to those of the count-based models. We provide scatter plots showing the relationship between call rate and the proportion of time spent vocalizing for each call-type category in Supplementary Figure S2, as well as model outputs in Supplementary Tables S7 and S8.

Supplementary Results S1: Inter-rater reliability

Across 5,733 paired segments, agreement was substantial ($\kappa$ = 0.846, z = 146, *p* <.001). Among the 5,187 segments detected by both raters, classification agreement was excellent ($\kappa$ = 0.957, z = 145, *p* <.001). Furthermore, on the level of broad functional categories (contact, alert, food), which was the level at which our analyses were conducted, kappa was 0.83, including unmatched segments (z = 94.4, *p* <.001) and 0.967 among matched segments (z = 96.2, *p* < .001), indicating that even when raters disagreed on the specific call type, they rarely annotated it as a call type from a different functional category.

Supplementary Results S2: Condition order trend analysis

Following the approach described in Supplementary Methods S2, we tested whether within-session condition presentation order had a significant effect on calling rates by adding polynomial trends (linear, quadratic, cubic) to model 1, 2, and 3A, restricted to the four randomized conditions (positions 3-6).

For total volubility (model 1), condition order did not significantly affect calling rates (χ^2^_(3)_ = 3.11, *p* = 0.375), and all substantive effects remained unchanged (condition: χ^2^_(3)_ = 76.33, *p*<.001, session: χ^2^_(1)_ = 19.87, *p* = 0.375; Table S6). Similarly, for contact call rates (model 2), condition order was non-significant (χ^2^_(3)_ = 3.82, *p* = 0.282), with session remaining the only significant predictor (χ^2^_(1)_ = 8.37, *p* = 0.004; Table S6).

For alertness calls across all conditions (model 3A), the condition order test reached significance (χ^2^_(3)_ = 8.81, *p* = 0.032). However, this was driven by a quadratic (U-shaped) trend rather than a linear decline, which is inconsistent with a monotonic habituation interpretation. All significant effects from the original analysis were preserved, including the interaction between condition and status (χ^2^_(3)_ = 32.27, *p* <.001), age (χ^2^_(1)_ = 5.46, *p* = 0.019), and session (χ^2^_(1)_ = 13.25, *p* <.001; Table S6).

Supplementary Results S3: Call rate vs. proportion of time spent vocalizing

We first examined the relationship between call rate and the proportion of time spent vocalizing at the level of individual observations across all calls together and all call-type categories separately, and additionally split contact calls into long-distance (phee-like calls and twitters) and short-distance (trill-like calls) ones (Figure S2). The two measures were strongly correlated across categories, with Pearson correlations ranging from 0.73 to 0.97 and Spearman correlations from 0.84 to 1.00 (all calls: r = 0.73, rs = 0.84; contact calls combined: r = 0.75, rs = 0.89; long-distance contact calls: r = 0.76, rs = 0.92; short-distance contact calls: r = 0.89, rs = 1.00; alertness calls: r = 0.97, rs = 1.00; food calls: r = 0.97, rs = 1.00). The highest correlations were observed for alertness, food, and short-distance contact calls, which are call types with relatively uniform syllable duration^3–5^, while all contact calls combined and long-distance contact calls alone showed somewhat lower but still substantial correlations, consistent with the larger range of syllable durations within these categories. These tight associations indicate that, despite differences in call duration across categories, the two measures capture closely overlapping aspects of vocal activity, with longer calls in some categories not systematically offsetting lower call counts.

Because models 1 and 2 cover the call-type categories with the lowest correlations and the largest variability in duration, we replicated these models using the proportion of time spent vocalizing as the response variable to verify that the conclusions drawn from call-rate analyses hold under this alternative measure.

For total volubility (model 1), the pattern of effects was highly consistent with the count-based model: condition remained a strong predictor (χ^2^_(5)_ = 52.55, *p* <.001), session showed a significant habituation effect (χ^2^_(1)_ = 27.00, *p* <.001), and neither age nor sex significantly predicted vocal output (Table S7). The planned contrasts likewise replicated the patterns observed in the count-based analyses (Table S8). One difference from the count-based analyses was that the effect of status class now reached marginal significance, with helpers spending more time vocalizing than breeders (odds ratio = 1.44, 95% CI [1.03; 2.02], χ^2^_(1)_ = 4.42, *p* = 0.035).

For contact calls (model 2), the pattern again closely mirrored the count-based analysis (Table S7): condition did not have a significant effect on the proportion of time spent vocalizing, with session remaining the only significant predictor (χ^2^_(1)_ = 8.15, *p* = 0.004) and status class showing a marginal trend (χ^2^_(1)_ = 3.59, *p* = 0.058). Consequently, none of the planned contrasts were significant, consistent with the call-rate model (Table S8). Together, these analyses indicate that the choice of vocal output measure does not substantively alter the conclusions of our study.
